# Delving into the *Bacillus cereus* group biosynthetic gene clusters cosmos: a comparative-genomics-based classification framework

**DOI:** 10.1101/2023.02.25.530005

**Authors:** Hadj Ahmed Belaouni, Amine Yekkour, Abdelghani Zitouni, Atika Meklat

## Abstract

**Background:** In this study, the *Bacillus* sp. strain BH32 (a plant-beneficial bacterial endophyte) and its closest non-type *Bacillus cereus* group strains were used to study the organization, conservation, and diversity of biosynthetic gene clusters (BGCs) among this group to propose a classification framework of gene cluster families (GCFs) among this intricate group. A dataset consisting of 17 genomes was used in this study. Genomes were annotated using PROKKA ver.1.14.5. The web tool antiSMASH ver. 5.1.2 was used to predict the BGCs profiles of each strain, with a total number of 198 BGCs. The comparison was made quantitatively based on a BGCs counts matrix comprising all the compared genomes and visualized using the Morpheus tool. The constitution, distribution, and evolutionary relationships of the detected BGCs were further analyzed using a manual approach based on a BLASTp analysis (using BRIG ver. 0.95); a phylogenetic analysis of the concatenated BGCs sequences to highlight the evolutionary relationships; and the conservation, distribution and the genomic co-linearity of the studied BGCs using Mauve aligner ver. 2.4.0. Finally, the BIG-SCAPE/CORASON automated pipeline was used as a complementary strategy to investigate the gene cluster families (GCFs) among the *B. cereus* group.

**Results:** Based on the manual approach, we identified BGCs conserved across the studied strains with very low variation and interesting singletons BGCs. Moreover, we highlighted the presence of two major BGCs synteny blocks (named “*synteny block* A” and “*synteny block* B”), each composed of conserved homologous BGCs among the *B. cereus* group. For the automatic approach, we identified 23 families among the different BGCs classes of the *B. cereus* group, named using a rational basis. The proposed manual and automatic approaches proved to be in harmony and complete each other, for the study of BGCs among the selected genomes.

**Conclusion:** Ultimately, we propose a framework for an expanding classification of the *B. cereus* group BGCs, based on a set of reference BGCs reported in this work.

## Background

The *Bacillus cereus* group of bacteria represents a homogeneous subdivision of the genus *Bacillus* with closely related phylogeny within the Firmicutes phylum [1, 2]. The bacteria constituting this group are Gram-positive, spore-forming, aerobic/facultative anaerobic, and rod-shaped with low-GC content [2]. Numerous bacteria related to *B. cereus* group were shown to produce several interesting compounds and enzymes, metabolize different kinds of pollutants, and promote the growth of both plants and animals when used as biostimulators [3–5]. The first described and most documented members of the group are *B. cereus*, *B. thuringiensis,* and *B. anthracis* [2]. Many bacteria classified as *B. cereus* are ubiquitous in the environment, with apparent soil origin, and present as commensal to intestines of insects or foodborne opportunistic pathogens often related to human poisoning [6, 7]. Other members of the group related to *B. thuringiensis* are insect pathogens widely used in agriculture for the biocontrol of insect pests [7, 8]; while *B. anthracis* is the causative agent of anthrax [2].

Traditionally, pathogenic potential and virulence characteristics permit organism differentiation within the group [1, 2, 7]: *B. cereus* by carrying biosynthetic gene cluster (BGC) for cereulide cytotoxin, plasmids carrying the crystal insecticidal genes for *B. thuringiensis* and presence of anthrax toxin and capsule genes for *B. anthracis* [2]. However, genetic experiments have exhibited a high level of synteny and protein similarity, with limited differences in gene content [1]. Conventional taxonomic markers, such as 16S and 23S rRNA genes, as well as whole-genome DNA hybridization seemed essentially identical, questioning the speciation of the group members [7]. Thus, extensive genomic similarities have contributed to the suggestion that *B. anthracis, B. cereus,* and *B. thuringiensis* are members of a single species, *B. cereus* sensu lato [1]. Moreover, the horizontal genetic transfer of plasmid and chromosome DNA among the strains of the *B. cereus* group has likely related to the diversity of this bacterial group, thus complicating the speciation [2].

In the present context of increased environmental screening with the generalization of whole genome sequencing that revealed the presence of newly identified recombinant forms [2], it is important to explore different approaches for understanding the evolution of the *B. cereus* group that may contribute to a more accurate characterization of these organisms. With the availability of a critical mass of compiled genomic data and the development of advanced computational tools for genome mining, one such approach consists of the bioinformatics exploration of an extensive range of potential biosynthetic gene clusters (BGCs) for the presence among the *B. cereus* group.

BGCs are a physical grouping of all the genes required to produce secondary metabolites, including pathway-specific regulatory genes [9]. BGCs mining is the process of identifying and characterizing these clusters to understand the biosynthesis of natural products and to discover new natural products and biosynthetic pathways [10]. BGCs are the source of many important natural products, such as antibiotics, anti-cancer compounds, and enzymes [11]. BGCs mining can lead to the discovery of new natural products with unique properties, and the development of new methods for natural product production [12]. BGCs can be used as a source of new enzymes and biosynthetic pathways that can be exploited for biotechnology applications such as the production of biofuels, fine chemicals, and enzymes for bioremediation [13]. Many natural products produced by BGCs have medicinal properties and can be used as leads for drug development. BGCs mining can lead to the discovery of new natural products with potential therapeutic applications [14]. BGCs are often horizontally transferred between bacteria [15]. To gain a genetic advantage, it is postulated that elements in BGCs are horizontally acquired across species for quick adaptation to a new environment [16]. Studying their evolution and distribution can provide insights into the evolution and ecology of bacteria [17].

With recent developments in next-generation sequencing and advancements in genome mining tools, it became possible to computationally identify thousands of BGCs and draw a global map of BGCs within a group of bacteria that allow us to systematically explore those of interest [9]. To overcome this challenge, researchers are increasingly using bioinformatics tools such as ClusterFinder [18], antiSMASH [19], and Big-scape [20], which can help automate the process of BGCs identification and classification. Additionally, efforts are being made to establish a standardized nomenclature for BGCs and to create a comprehensive database of BGCs, which lead to the establishment of the MIBiG database [21, 22] which can help facilitate data sharing and comparison among researchers.

Big-scape is a bioinformatics tool that can be used to study BGCs in bacteria using an automated streamlined pipeline [20]. It enables researchers to identify, classify, and compare BGCs across different bacterial strains and can be used to infer the biosynthetic pathways and natural products associated with each BGC, providing valuable insights into the evolution and adaptation of bacteria to different environments, as well as the discovery of new natural products and biosynthetic pathways. Automated bioinformatics pipelines and manual bioinformatics are both useful methods for analyzing biological data, but they have different advantages and disadvantages. Automated pipelines are more efficient and can be run by researchers with minimal bioinformatics experience [23], while manual bioinformatics is more flexible and customizable, but requires more expertise [24]. The choice of which method to use will depend on the specific research question, the amount and type of data, and the available resources.

Based on the known biosynthesis pathways potentially involved in the production of specialized metabolites by *Bacillus* and closely related species, BGC predictions rely on the development of bioinformatics tools and algorithms design to search for conserved motifs of specific pathways; including peptide synthetases (NRPSs), polyketide synthases (PKSs), and ribosomally synthesized and post-translationally modified peptides (RiPPs) pathways [25]

Despite BGCs being important among the *Bacillus cereus* group, there is currently limited data on their conservation across the different strains of this group. Further research is required to better understand the diversity and distribution of BGCs among this group. Nevertheless, highlighting the synteny of BGCs in bacteria is one challenging yet beneficial task.

Synteny refers to the preservation of gene order and chromosomal location among different organisms [26]. One benefit of studying this phenomenon is to guide the discovery and characterization of new natural products produced by these bacteria. By identifying the synteny of BGCs among different strains, researchers can infer the presence of similar BGCs in other strains, and can then use this information to guide their search for new natural products. Another benefit is that it can provide insight into the evolutionary relationships among different taxa. The conservation of BGCs in terms of chromosomal location and gene organization among different strains can be used to infer evolutionary relationships among these strains, and can also help to understand how these bacteria have adapted to different environments over time.

Advances in sequencing technologies, high-throughput screening techniques, and improved computational methods have led to a rapidly increasing number of BGCs being identified. This has created a growing need for effective and standardized methods for BGC classification [12]. There have been several efforts to establish a unified system for BGC classification, such as the antiSMASH platform, but the field is still in its early stages and much work remains to be done to develop a comprehensive and widely-accepted classification system [19].

Moreover, during a screening for potentially beneficial endophytic bacteria, the strain *Bacillus* sp. BH32, which belongs to the *B. cereus* group, was isolated from *Atriplex halimus L*., a halophyte sampled from a continental hypersaline region (Sebkha) in Djelfa province, Algeria. The strain, which was proven to help tomato and wheat seedlings tolerate salt stress at various levels, was consecutively genome analyzed to determine putative mechanisms involved in salt tolerance and plant promotion [27], and thus, is part of the genome dataset of this study, along with its closely-related strains.

In the present study, we investigated the BGCs of the *Bacillus* sp. strain BH32 at a genomic level along with its closest non-type strains to explore the conservation and putative evolution patterns of BGCs among the *Bacillus cereus* group, and to highlight singletons. Based on a combined strategy (manual and automatic), we aimed to establish the basis of a rational classification of BGCs among the *B. cereus* group.

## Methods

### 1. Presenting the dataset

The genomic dataset is composed of *Bacillus* sp. BH32 genomes and its closest non-type strains genomes; all part of the *Bacillus cereus* group. *Bacillus* sp. BH32 is a beneficial endophyte, isolated during a previous study from *Atriplex halimus* L., a halophyte sampled from an Algerian continental Sebkha from the province of Djelfa. This strain was proven to help tomato and wheat seedlings tolerate salt stress at various levels [27]. The selection of the closest non-type strains of *Bacillus* sp. BH32 was performed with BLASTn 2.10.0+ [28], using the whole genome of *Bacillus* sp. BH32 as a query against the NCBI’s “Complete prokaryotic genomes” database (https://blast.ncbi.nlm.nih.gov/Blast.cgi, accessed on 02-04-2020), targeting the first most significant 16 hits (high-quality complete genomes only) **(table 1)**.

**Table 1.** Dataset for the comparative analysis

| Preferred name | Deposit | Base pairs | % G+C | Proteins | Bioproject accession | Biosample accession | Assembly accession |
| --- | --- | --- | --- | --- | --- | --- | --- |
| *Bacillus wiedmannii* | PL1 | 5,309,441 | 35.26 | 5272 | PRJDB9286 | SAMD00204526 | GCA_011405335.1 |
| *Bacillus* sp. | BH32 | 5,661,597 | 34.86 | 5682 | PRJNA523918 | SAMN10992522 | GCF_004367825.2 |
| *Bacillus cereus* | JHU | 5,323,903 | 35.26 | 5489 | PRJNA591929 | SAMN13391868 | GCA_009738575.1 |
| *Bacillus cereus* | G9842 | 5,387,334 | 35.26 | 5379 | PRJNA224116 | SAMN02604060 | GCF_000021305.1 |
| *Bacillus thuringiensis* | HD-789 | 5,495,278 | 35.26 | 5550 | PRJNA171844 | SAMN02603564 | GCA_000292705.1 |
| *Bacillus bombysepticus* | Wang | 5,295,783 | 35.25 | 5265 | PRJNA242213 | SAMN02691825 | GCF_000831065.1 |
| *Bacillus thuringiensis* | HD1002 | 5,491,311 | 35.25 | 5548 | PRJNA236049 | SAMN03010437 | GCF_000835025.1 |
| *Bacillus thuringiensis* serovar *galleriae* | 4G5 | 5,701,188 | 35.29 | 5770 | PRJNA224116 | SAMN03074947 | GCF_000803665.1 |
| *Bacillus thuringiensis* serovar *morrisoni* | BGSC 4AA1 | 5,652,292 | 35.33 | 5693 | PRJNA224116 | SAMN03274640 | GCF_000940785.1 |
| *Bacillus cereus* | A1 | 5,352,307 | 35.28 | 5273 | PRJNA242371 | SAMN02693464 | GCA_000635895.2 |
| *Bacillus thuringiensis* | YGd22-03 | 5,420,545 | 35.23 | 5455 | PRJNA224116 | SAMN06197294 | GCF_002184245.1 |
| *Bacillus thuringiensis* | L7601 | 5,790,408 | 35.18 | 5846 | PRJNA224116 | SAMN06473083 | GCF_002025105.1 |
| *Bacillus thuringiensis* | c25 | 5,334,660 | 35.32 | 5300 | PRJNA388287 | SAMN07317425 | GCA_002222555.1 |
| *Bacillus cereus* | FORC087 | 5,271,204 | 35.27 | 5272 | PRJNA470818 | SAMN09111977 | GCA_006384875.1 |
| *Bacillus cereus* | ZB201708 | 5,466,652 | 35.22 | 5423 | PRJNA480899 | SAMN09652006 | GCA_004006495.1 |
| *Bacillus thuringiensis* | BT-59 | 5,500,615 | 35.26 | 5549 | PRJNA534189 | SAMN11479299 | GCA_005155285.1 |
| *Bacillus thuringiensis* | JW-1 | 5,500,376 | 35.26 | 5552 | PRJNA574201 | SAMN12860338 | GCA_009025915.1 |

### 2. BGCs counts and distribution

BGCs from the dataset were predicted using antiSMASH ver. 5.1.2 [19]. BGCs counts were compared quantitatively, after preparing a BGCs-types/counts matrix comprising all the compared genomes, and visualized using Morpheus (https://software.broadinstitute.org/morpheus/). The generated heatmap showed the counts of BGCs (cyan-white-fuchsia, from 0 to 5) by type, and their distribution among the strains (dendrograms were generated by hierarchical clustering using one minus Spearman rank correlation) **(Fig. 1).**

### 3. BGCs BLASTp comparisons

A BLASTp (ver. 2.9.0+) comparison of *Bacillus* sp. BH32 BGCs against BGCs of the closest non-type strains was done, using the BGCs amino-acid sequences (for each genome, a multifasta file was used, containing the amino acid sequences of all the genes constituting the whole predicted BGCs, prepared from the GenBank records generated by antiSMASH), using BRIG ver. 0.95 [29]. The reference ring constitutes the detected BGCs in *Bacillus* sp. BH32, while each ring represents BGCs of each genome of the closest non-type strains (the color shades represent sequence identity, the greyer it gets; the lower the percentage identity). The 3 outermost rings (from inside to outside) represent: genes composing BGCs in *Bacillus* sp. BH32 according to their respective location in the genome; unique genes (sequence identity < 30% with 100% of the strains) and rare genes (sequence identity < 30% with 80% the strains) with their locus tag IDs; and the corresponding regions/BGCs types (last ring) as detected by antiSMASH **(Fig. 2)**. The annotation from antiSMASH and PROKKA of each gene was retrieved using their corresponding locus tag IDs.

### 4. BGCs gains, losses, and rearrangements

#### Manual approach

First, fasta files were prepared; each containing concatenated amino acid sequences of all BGCs for each strain (with conserved order of occurrence in the respective genome). Sequences were aligned using MAFFT ver. 7.221.3 [30]. An ML tree was inferred from the aligned sequences by using the Maximum Likelihood method based on the JTT matrix-based model [31]. The tree harboring the highest log likelihood (-248721.7967) was selected. The percentage of trees in which the associated sequences clustered together is displayed next to the branches (branch-support values). Initial tree(s) for the heuristic search were automatically obtained based on Neighbor-Join and BioNJ algorithms applied to a matrix of pairwise distances (estimated by a JTT model), and then selecting the topology with the highest log likelihood value. The tree was drawn to scale, with branch lengths reflecting the number of substitutions per site. The analysis involved 17 amino acid sequences (full record of BGCs from each strain). All positions containing gaps and missing data were eliminated. There were a total of 28898 positions in the final dataset. Evolutionary analyses were conducted in MEGA7 [32]. The final tree was drawn using Adobe Illustrator CC 18.1.1 **(Fig. 3 a).**

BGCs synteny among the strains was investigated with MAUVE ver. 2.4.0 [33–37], using the alignment of concatenated Genbank sequences of the BGCs of each strain, according to their order of appearance in the respective genome, where each row shows the conservation and orientation of BGCs in the corresponding strain in the ML tree after alignment. *Bacillus thuringiensis* c25 was set as a reference, according to the phylogenetic analysis (most distant). Homologous segments indicating orthologous clusters (from MAUVE alignment diagram) with a locally collinear block (LCB) weight ≥ 773 were confirmed by annotation. BGCs relative orientation and order conservation were highlighted **(Fig. 3 b).**

#### BIGSCAPE automatic approach and GCFs nomenclature proposal

The selected antiSMASH profiles of the *B. cereus* genomes were used to identify the gene cluster families GCFs using BiG-SCAPE v.1.1.0. with default parameters [20]. The output of the BIG-SCAPE was exploited to propose a classification of GCFs among the *B. cereus* group, based on the respective BGCs classes/types, as organized by the unsupervised Machine Learning clustering approach employed by BIG-SCAPE. Since the BIG-SCAPE output consists of BGCs grouped into GCFs with arbitrary codes, we propose a systematic and grounded nomenclature of the highlighted families. In our proposal, families names are composed of: a **“**BC” prefix that stands for *Bacillus cereus* (group) (“BCS” in case of singletons), followed by the type (e.g. bacteriocin), then an increasing number that should follow the order of detection/characterization during this study (or future studies). The product’s name should appear instead of merely the type, in case the cluster product is known (from the MIBIG database). The proposed reference BGCs are based on ‘*exemplars’*, according to the affinity propagation clustering approach used by BIG-SCAPE. Harmony between the manual and automatic approaches was assessed manually based on the correspondence between the BGCs region codes as given by antiSMASH and those displayed among the BIG-SCAPE output. The similarity networks were arranged to fit their corresponding figures, while the original BIG-SCAPE output, the used dataset (antiSMASH profiles), and the proposed reference BGCs files (included in the antiSMASH profiles) are available as a supplementary material. Final figures of the GCFs classes distribution (Fig. 5-8) were composed under Adobe Illustrator CC 18.1.1.

## Results

### 1. BGCs distribution and counts in the *B. cereus* group

The BGCs were first compared in terms of distribution and counts among the compared genomes **(Fig. 1)**. All the strains have similar counts of NARPS-like, LAP-bacteriocin, siderophore, and betalacton BGCs (one for each type). *B. cereus* JHU has the highest number of bacteriocin BGCs (4), followed by *B. thuringiensis* L-76-01 and B. thuringiensis c25 (3 each), while the other strains have 2 (except *Bacillus* sp. BH32, only one bacteriocin BGC was detected). *B. cereus* ZB201708 is the only strain where antiSMASH failed to detect a terpene BGC (all the other strains have one). *B. cereus* ZB201708, *B. thuringiensis* YGd 22-03, and *B. thrungiensis* c25 are the only strains to have a lanthipeptid BGC (one for each). NRPS BGC-type has the highest counts among the selected strains, ranging from 3 to 5. *Bacillus cereus* A1, *Bacillus bombysepticus* Wang, *Bacillus thuringiensis* c25 and *Bacillus cereus* FORC087 have 3 NRPS BGCs ; *Bacillus* sp. BH32, *Bacillus wiedmannii* PL1, *Bacillus thuringiensis* YGd22-03 and *Bacillus thuringiensis* serovar galleriae 4G5 have 4 ; and the remaining strains have 5.

The strains *Bacillus thuringiensis* serovar morrisoni BGS, *Bacillus cereus* G9842, *Bacillus thuringiensis* BT59, *Bacillus thuringiensis* HD1002, *Bacillus thuringiensis* JW-1, and Bacillus *thuringiensis* HD 789 have the same BGC counts (NRPS = 5, NRPS-like = 1, LAP/bacteriocin = 1, bacteriocin = 2, terpene = 1, betalactone = 1, siderophore = 1, lanthipeptide = 0). Same thing for *Bacillus wiedmannii* PL1 and *Bacillus thuringiensis* serovar galleriae 4G5 (NRPS = 4, NRPS-like = 1, LAP/bacteriocin = 1, bacteriocin = 2, terpene = 1, betalactone = 1, siderophore = 1, lanthipeptide = 0). *Bacillus cereus* A1, *Bacillus bombysepticus* Wang and *Bacillus cereus* FORC087 have the same profile as well (NRPS = 3, NRPS-like = 1, LAP/bacteriocin = 1, bacteriocin = 2, terpene = 1, betalactone = 1, siderophore = 1, lanthipeptide = 0). The remaining strains have different BGC count profiles **(Fig. 1)**.

**Fig. 1.**
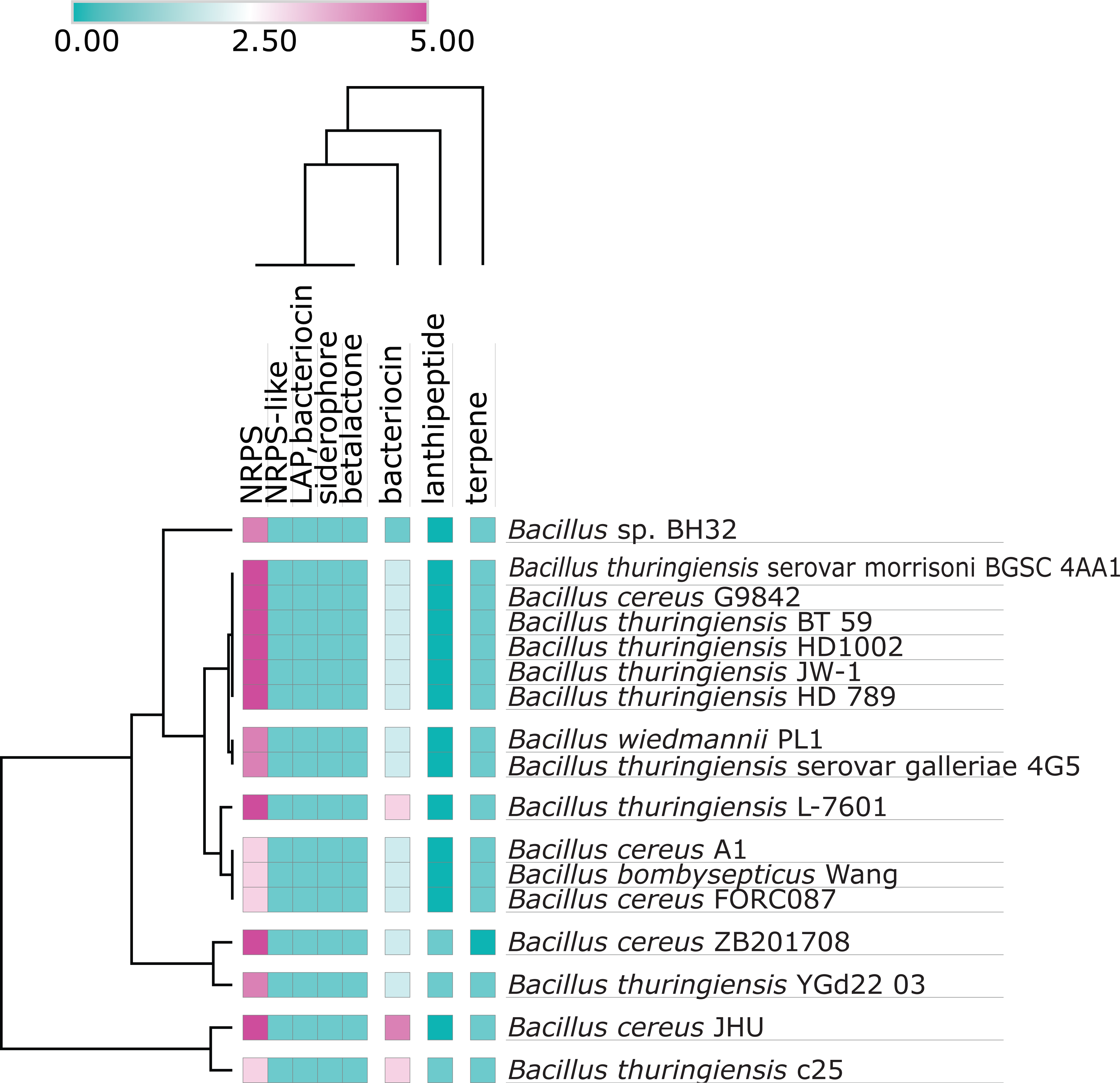
Distribution of BGCs classes and counts. The heatmap shows the number of BGCs by type (from 0 to 5), and their distribution among the strains (dendrograms were generated by hierarchical clustering using *one minus Spearman* rank correlation)

### 2. Amino acid sequence conservation of the *B. cereus* group BGCs

The BLASTp comparison of BGCs from *Bacillus* sp. BH32 vs. its closest non-type strains **(Fig. 2)** shows heterogeneous content in terms of predicted protein sequences from detected BGCs. The most conserved BGC seems to be the terpene BGC (region 2.1), while the less conserved one seems to be the NRPS BGC (region 2.2).

Although the antiSMASH failed to detect the terpene biosynthetic genes, they seem to be present in the BLASTp output of *Bacillus cereus* ZB201708 (14^th^ ring, from the inner ring to the outer ring), which could be due to assembly concerns, or else, the constituting genes of this terpene cluster might be distributed in other BGCs from *Bacillus cereus* ZB201708.

NRPS BGC (region 2.2) from *Bacillus* sp. BH32 has very low sequence identity when compared to its counterparts in *B. bombysepticus* Wang, *B. cereus* A1 and *B. thuringiensis* c25 (<30%). Considering that the other NRPS BGCs from *Bacillus* sp. BH32 (regions 6.1, 8.1, 12.1, and 15.1) have higher sequence identity % with the 3 corresponding BGCs of the aforementioned strains, which confirms the absence of a counterpart of the NRPS BGC from region 2.2 in these strains as suggested by the counts (5 NRPS BGCs in *Bacillus*. sp. BH32 against only 4 in *B. bombysepticus* Wang, *B. cereus* A1, and *B. thuringiensis* c25). This cluster has two rare genes (identity bellow 30% for 80% of the strains): ctg2_300 = unknown/hypothetical protein; ctg2_321 = unknown / IS200/IS605 family transposase ISAsp8.

The siderophore BGC (region 3.1) encoding petrobactin (100%) and the LAP/bacteriocin BGC (region 5.1, unknown encoded product, closest BGCs with overall gene sequence identity >70% from strains *Bacillus cereus* ZB201708, *Bacillus cereus* JHU and *Bacillus thuringiensis* L-7601) seem to be well conserved, with a slight variation on the composing genes. The NRPS BGC (region 6.1) is another example of a well-conserved cluster across the compared strains (closest known cluster = bacillibactine, 46%).

The NRPS-like BGC (region 8.1) is present among all the strains. Nevertheless, *Bacillus* sp. BH32 has one unique gene (sequence identity bellow 30% for 100% of the strains) in this region (ctg8_81 = unknown/hypothetical protein), and a rare one (ctg8_90 = unknown/hypothetical protein).

The NRPS BGC (region 12.1) shows variation in its first 19 genes (∼ 1/3 of the total BGC), including one unique gene (ctg12_31 = unknown / Spore germination protein B1), and one rare gene (ctg12_35 = biosynthetic-additional; (smcogs) SMCOG1012:4’-phosphopantetheinyl transferase / 4’-phosphopantetheinyl transferase Sfp).

The NRPS BGC (region 15.1) from *Bacillus* sp. BH32 has one unique gene (ctg15_85 = biosynthetic-additional; (smcogs) SMCOG1028: crotonyl-CoA reductase - alcohol dehydrogenase / Zinc-type alcohol dehydrogenase-like protein). This NRPS BGC doesn’t have a counterpart in *Bacillus cereus* FORC087, confirming the BGCs count (5 NRPS BGCs in *Bacillus* sp. BH32 for only 4 in *Bacillus cereus* FORC087).

The bacteriocin BGC (region 24.1) from *Bacillus* sp. BH32 is another example of a well-conserved cluster across its closely related non-type strains.

Finally, the betalacton BGC (region 27.1) from *Bacillus* sp. BH32 (fengycin, 40%) arbores two unique genes (ctg27_13 = unknown/hypothetical protein; ctg27_20 = unknown/hypothetical protein). Globally, all BGCs from *Bacillus* sp. BH32 are present in at least 13 of its closest strains (>81%), either very well-conserved (regions) or less conserved, with some rare/unique genes **(Fig. 2)**.

**Fig. 2.**
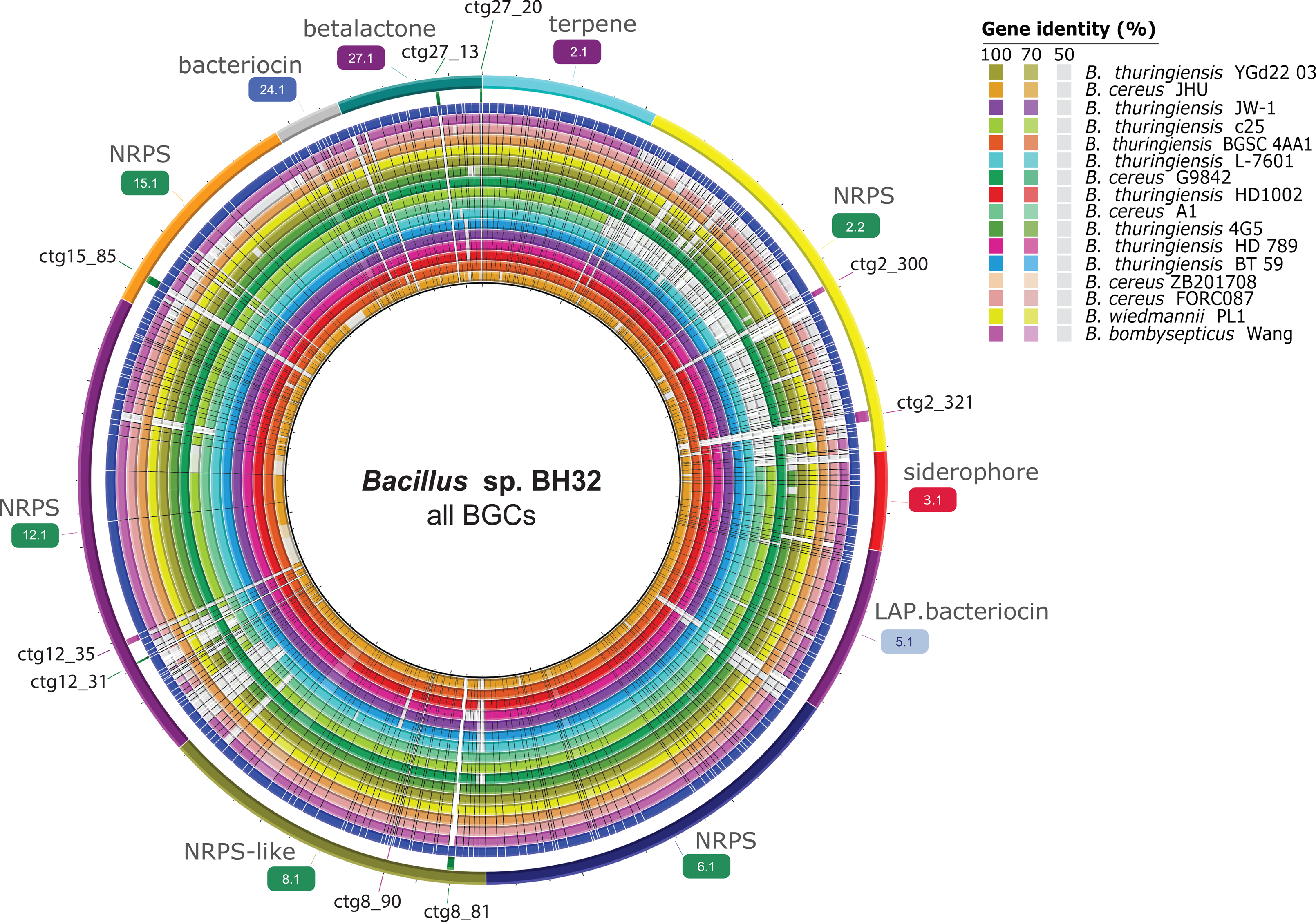
BLASTp comparisons of *Bacillus* sp. BH32 BGCs against BGCs of the closest non-type strains. The reference ring constitutes the detected BGCs in *Bacillus* sp. BH32 (translated to amino acid sequence), while individual rings represent BGCs of the closest non-type strains (the color represents sequence identity on a sliding scale, the greyer it gets; the lower the percentage identity). The 3 outermost rings depict (from inside to outside): genes composing BGCs in *Bacillus* sp. BH32 (in blue); *unique genes* (sequence identity bellow 30% for 100% of the strains, in green) and *rare genes* (sequence identity bellow 30% for 80% of the strains, in light velvet) with their locus tag IDs; and the *regions/BGCs types* (as detected by antiSMASH). Unique and rare genes have the following annotation (from antiSMASH / prokka, with the same location, or overlapping locations): ctg8_81 = unknown/hypothetical protein; ctg12_31 = unknown / Spore germination protein B1; ctg 15_85 = biosynthetic-additional (smcogs) SMCOG1028:crotonyl-CoA reductase - alcohol dehydrogenase / Zinc-type alcohol dehydrogenase-like protein; ctg27_13 = unknown/hypothetical protein; ctg27_20 = unknown/hypothetical protein; ctg 2_300 = unknown/hypothetical protein; ctg2_321 = unknown / IS200/IS605 family transposase ISAsp8 ; ctg8_90 = unknown / hypothetical protein; ctg 1 2_35 = biosynthetic-additional (smcogs) SMCOG1012:4’-phosphopantetheinyl transferase / 4’-phosphopantetheinyl transferase Sfp

### 3. Manual approach for *B. cereus* group BGCs synteny

The ML tree based on BGCs amino acids sequences **(Fig. 3 a)** shows 4 clades:

Clade I: *B. thuringeinsis c25, B. cereus* JHU, *B. thuringeinsis* HD1002 and *B. thuringeinsis* BT-59; Clade II: *Bacillus* sp. BH32; Clade III: *B. wiedmannii* PL1, *B. cereus* FORC087, *B. thuringiensis* serovar galleriae 4G5, *B. bombysepticus* Wang, *B. cereus* A1 and *B. thuringiensis* YGd22-03; Clade IV: B. thuringiensis HD 789, B. thuringiensis JW1, *B. thuringiensis* L-7601, *B. cereus* G9842, *B. thuringiensis* serovar morrisoni BGS and *B. cereus* ZB201708.

From the ML tree **(Fig. 3 a)**, *B. thuringiensis* c25 has been considered as a reference for BGCs conservation (the most distant genome among the compared strains). *Bacillus* sp. BH32 which usually clusters with *B. cereus* ZB201708 and *B. thuringiensis* serovar morrisoni BGSC 4AA1 (GBDP and ANI analysis), doesn’t seem to be close to these strains when it comes to its BGCs, showing thus more complexity among this intricate group, in terms of BGCs.

The analysis of the BGCs conservation is shown in **(Fig. 3 b)**. Here, BGCs are mentioned according to their attributed BGC tag number **(Fig. 3 b** legends). Considering *B. thuringiensis* c25 as a reference, we observe that the orthologous BGCs group that seems to be best conserved in order and appearance is the one composed of BGC3 (bacteriocin: unknown), BGC4 (bacteriocin: unknown), BGC5 (betalacton, fengycin), BGC6 (NRPS: gramicidin in *B. cereus* JHU; nostopeptolide A2 in *B. thuringiensis* serovar galleriae 4G5; unknown in the remaining strains), BGC7 (NRPS: bacillibactin), BGC8 (siderophore: petrobactin) and BGC9 (linear azol(in)e-containing peptides “LAP” bacteriocin). This group was named “*synteny block A”*, which appears in this order in all strains except *B. wiedmannii*, where BGC3 appears to be in the beginning, unusually separated from the other BGCs.

The second conserved group is composed of BGC1 (terpene: molybdenum co-factor) and BGC2 (NRPS: polyoxypeptin, except for *B. thringiensis* c25, unknown), named “*synteny block B”*. BGC10 (bacteriocin: unknown) is only present in *B. thuringiensis* C25 (thus, will not be mentioned here again).

In *B. cereus* JHU, BGC1 and BGC2 (*synteny block B*) are still in the same order, as well as for the BGC3, BGC4, BGC5, BGC6, BGC7, BGC8 and BGC9 (*synteny block A*). Meanwhile, 4 new BGCs appeared: BGC13 (bacteriocin: unknown) and BGC14 (bacteriocin: unknown) both in the beginning; BGC15 (NRPS-like 2: unknown) between the two synteny blocks A and B; and BGC16 (NRPS-Polyketide: chejuenolide A / chejuenolide B) at the end. The BGC11 (NRPS-like 1: unknown) moved to the beginning, right before the *synteny block B*. BGC12 (lanthipeptid: cerecidin / cerecidin A1 / cerecidin A2 / cerecidin A3 / cerecidin A4 / cerecidin A5 / cerecidin A6 / cerecidin A7) is lost.

BGC13 and BGC14 (both unknown bacteriocins) are only found in *B. cereus* JHU.

*As for B. cereus* JHU, *B*. *thuringiensis* HD-1002 and *B. thuringiensis* BT-59 have the two synteny blocks with the BGC15 in between. BGC16 is at the right end of the *synteny block B*. BGC12, BGC13, and BGC14 are lost and the BGC11 jumped to the end.

Since *Bacillus* sp. BH32 genome doesn’t constitute a complete one, the observations for the BGCs conservation for this strain concern only their presence, not their order (the use of a reference genome for the reordering of the contigs doesn’t take into count the possible genomic rearrangements, and thus biases the synteny analysis, so we kept the contigs as they resulted from the assembly, from the largest to the smallest one).

In *Bacillus* sp. BH32, the BGC4 (bacteriocin with no closest known cluster) seems to be missing, while conserved across all the analyzed genomes of its closely related non-type strains, suggesting this absence due to the incompleteness of the genome, and should be found once the genomic gaps filled. The remaining BGCs from *synteny blocks A* and *B* are all present in *Bacillus* sp. BH32.

In the BGCs clades III and IV, all BGCs are inversely oriented in comparison with the strains of the first clade (including *B. thuringiensis* C25). In *B. wiedmannii,* BGC11 jumped between *synteny blocks A* and *B*, while BGC3 and BGC15 are right before the *synteny block B*. This strain doesn’t have the BGC12.

*B. cereus* FORC087 has the same homologous BGCs from 1 to 9 (inversely oriented), with BGC11 in the beginning. BGC15 and BGC16 are absent. *B. thuringiensis* serovar galleriae 4G5 has *synteny blocks A* and *B*, and the BGC15 reappears next to BGC2 from *synteny block B*. As for *B. cereus* FORC087, BGC 11 is still in the beginning.

For *B. bombysepticus* Wang, BGC2 is missing. Despite the BGC15 (NRPS-like: unknown) bears homologous segments linking it with the BGC2 (NRPS: polyoxypeptin), after manually checking the annotation, it confirmed the assignment to its closest orthologous BGC, the BGC15. The BGC1 is there, at the end. BGC11 is still in the beginning, before the *synteny block A*.

*B. cereus* A1 has an arrangement similar to *B. bombysepticus* Wang, except for BGC11 jumping to the end. For *B. thuringiensis* YGd22-03, BGC2 reappears, in the *synteny block B*, while the BGC12 lost since *B. thuringiensis* c25 reappears here.

For clade IV, the *synteny blocks A*, *B,* and BGC15 hold the same position in all its strains. In *B. thuringiensis* HD-789, BGC11 is in the end, next to the *synteny block B*. BGC16 reappears, to stand in the beginning.

*B. thuringiensis* HD-789*, B. thuringiensis JW-1, B. thuringiensis* serovar morrisoni BGSC 4AA1 *B. cereus* G9842*, B. thuringiensis L76-01* and *B. cereus* ZB201708 have the BGC11 jumping again to the front position, followed by the BGC16, except *B. cereus* ZB201708 where the appearing BGC18 (lanthipeptid: surfactin) appeared between BGC11 and BGC16; and the *synteny blocks A* and *B* with BGC15 in between. *B. thuringiensis L76-01* and *B. cereus* ZB201708 are characterized with a new BGC for each one: BGC17 (bacteriocin: unknown) and BGC19 (bacteriocin: unknown) respectively, at the end.

BGCs of the *synteny blocks A* and *B* being consistently present in all the strains, will naturally tend to occur in chromosomic locations among the analyzed genomes. BGC11 (NRPS-like 1: unknown) is present across all strains, at various positions, suggesting its possible jumping between chromosomic and plasmidic locations.

BGC15 (NRPS-like 2: unknown) is largely present (except in *B. thuringiensis* c25 and *B. cereus* FORC087), and is located most of the time between synteny blocks A and B, suggesting a more likely chromosomic location as well. BGC12 (lanthipeptid: cerecidin / cerecidin A1 / cerecidin A2 / cerecidin A3 / cerecidin A4 / cerecidin A5 / cerecidin A6 / cerecidin A7), when present, is next to BGC11.

The similarity plots in the homologous segments are confirming the BLASTp results from the BRIG ring generator **(Fig. 2)**. The present synteny analysis can give direction to the prediction of the presence of certain BGCs, as we showed that they are usually conserved in presence and order. It can also be used to detect the evolution and distribution of homologous BGCs across the complex *B. cereus* group. The observed BGCs positions should be further investigated, in their genomic context, by studying synteny not only for BGCs, but extending the analysis to the whole genomic content.

**Fig. 3.**
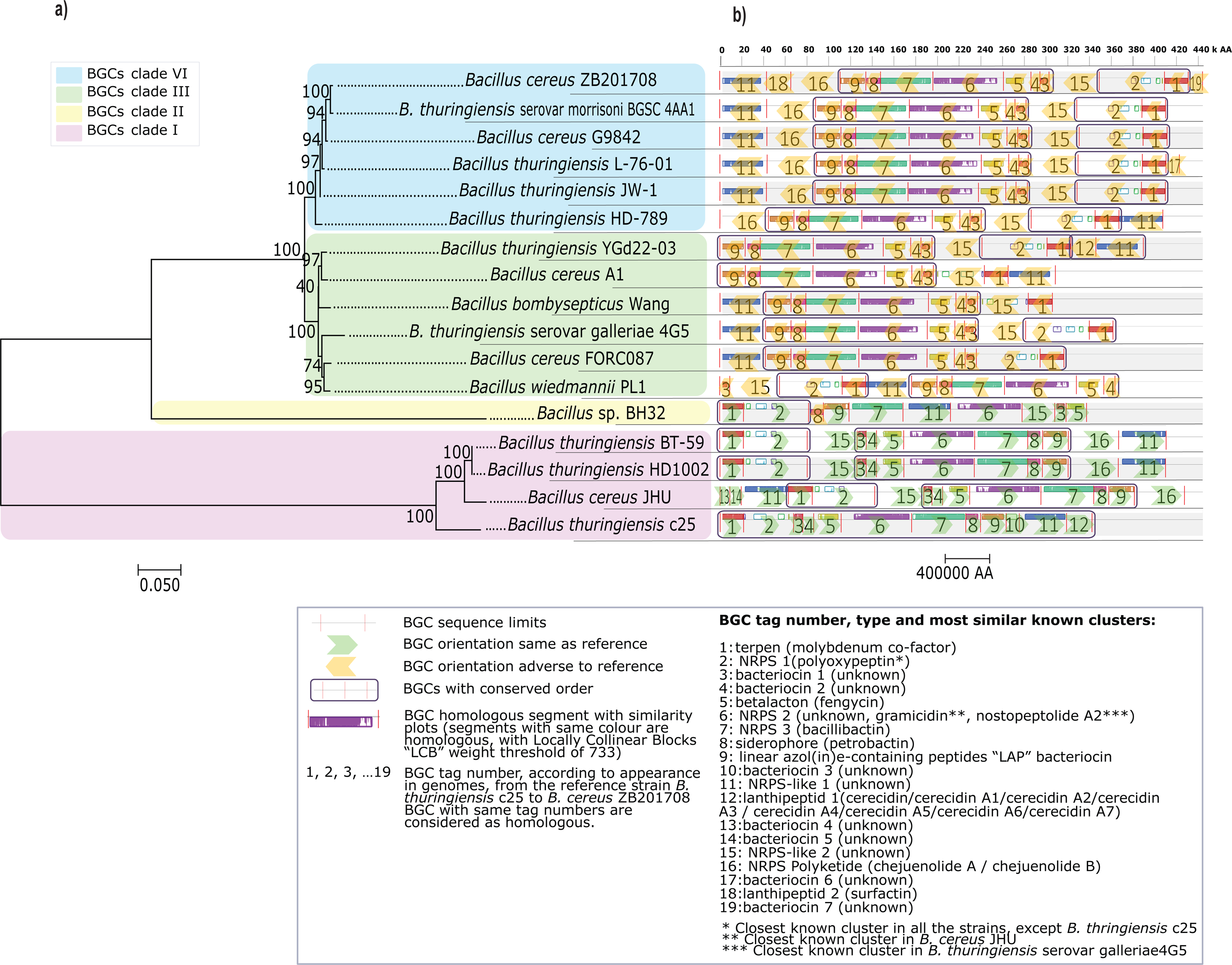
BGCs phylogeny and conservation. **a)** Maximum Likelihood (ML) phylogenetic tree generated from concatenated BGCs amino acid sequences. Numbers on branches represent bootstrap values (average value: 92.21%). The tree is drawn to scale, with branch lengths measured in the number of substitutions per site. **b)** BGCs conservation among the strains (MAUVE alignment of concatenated genebank sequences of the BGCs of each strain). Each row represents the conservation and orientation of BGCs of the corresponding strain in the ML tree (left) after alignment, in comparison to *Bacillus thuringiensis* c25 (bottom row, set as reference). Red bars represent BGC sequence limits. Colored blocks refer to homologous segments with similarity plots (from MAUVE alignment diagram), with a *locally collinear block* (LCB) weight ≥ 773. Each arrow symbolizes the BGC relative orientation with its tag number. BGCs with conserved order are framed in purple. BGC tag number, type, and most similar known clusters are shown on legends. The distance scale is shown under the alignment diagram (in amino acids)

### 4. BIG-SCAPE results

The BIG-SCAPE pipeline was used as well, in line with the manual approach, as a complementary strategy to investigate the gene cluster families (GCFs) among the *B. cereus* group.

There was a total of 198 BGCs, organized into 4 main classes (Fig. 4, table 2):

• NRPS: consisting of a total of 89 BGCs, it is the biggest class representing 44.94% of the total BGCs;
• RiPPs: 58 BGCs fall into this class, representing 29.29% of the sum of the analyzed BGCs;
• Terpenes: represent 5.58% of the total BGCs (with 17 BGCs identified as coding for a terpene);
• Other (remaining classes): composed of 34 BGCs, representing 17.17% of the studied BGCs.

**Table 2.**
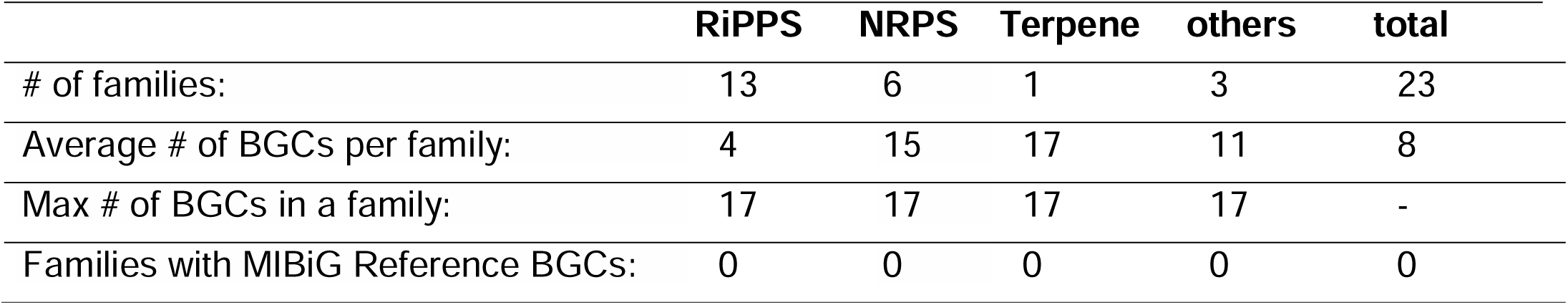
BGC classes/families overview

|  | RiPPS | NRPS | Terpene | others | total |
| --- | --- | --- | --- | --- | --- |
| # of families: | 13 | 6 | 1 | 3 | 23 |
| Average # of BGCs per family: | 4 | 15 | 17 | 11 | 8 |
| Max # of BGCs in a family: | 17 | 17 | 17 | 17 | - |
| Families with MIBiG Reference BGCs: | 0 | 0 | 0 | 0 | 0 |

We suggest a framework for an expanding classification of the *B. cereus* group BGCs, based on a set of reference BGCs described in this work (tables 3-4; supplementary figures S1, S2, S3, and S4), as anchoring points to affiliate unknown query BGCs to the proposed families/clans accordingly. Such reference BGCs should be included in future attempts to assign a set of BGCs (from genomes belonging to the *B. cereus* group) using the BIG-SCAPE pipeline into one of the proposed families in the current proposal. Moreover, the same strategy can be reproduced for other bacterial groups.

**Table 3.**
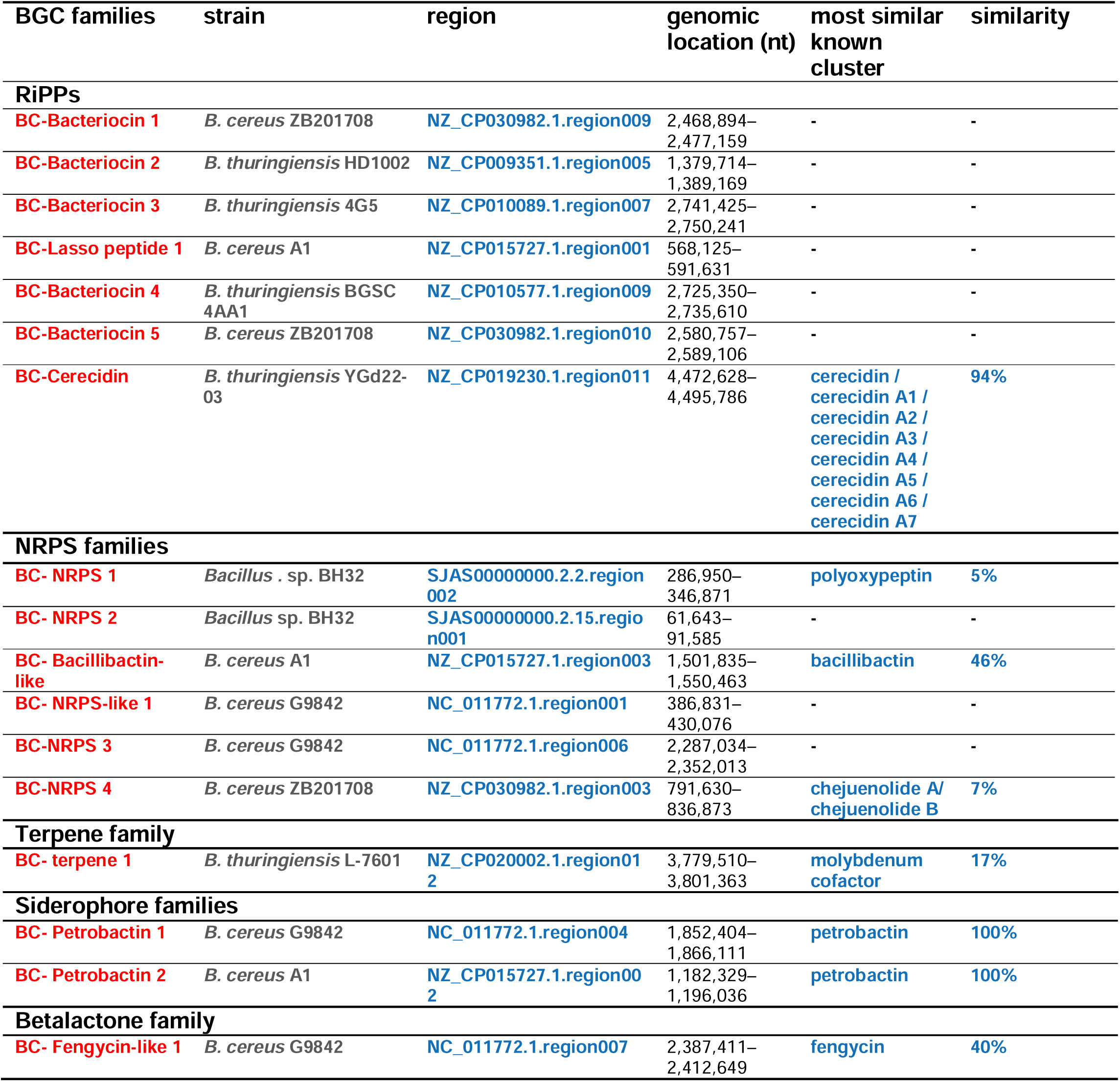
Reference BGCs per class/family.

| BGC families | strain | region | genomic location (nt) | most similar known cluster | similarity |
| --- | --- | --- | --- | --- | --- |
| **RiPPs** |  |  |  |  |  |
| **BC-Bacteriocin 1** | *B. cereus* ZB201708 | NZ_CP030982.1.region009 | 2,468,894–2,477,159 | - | - |
| **BC-Bacteriocin 2** | *B. thuringiensis* HD1002 | NZ_CP009351.1.region005 | 1,379,714–1,389,169 | - | - |
| **BC-Bacteriocin 3** | *B. thuringiensis* 4G5 | NZ_CP010089.1.region007 | 2,741,425–2,750,241 | - | - |
| **BC-Lasso peptide 1** | *B. cereus* A1 | NZ_CP015727.1.region001 | 568,125–591,631 | - | - |
| **BC-Bacteriocin 4** | *B. thuringiensis* BGSC 4AA1 | NZ_CP010577.1.region009 | 2,725,350–2,735,610 | - | - |
| **BC-Bacteriocin 5** | *B. cereus* ZB201708 | NZ_CP030982.1.region010 | 2,580,757–2,589,106 | - | - |
| **BC-Cerecidin** | *B. thuringiensis* YGd22-03 | NZ_CP019230.1.region011 | 4,472,628–4,495,786 | cerecidin / cerecidin A1 / cerecidin A2 / cerecidin A3 / cerecidin A4 / cerecidin A5 / cerecidin A6 / cerecidin A7 | **94%** |
| **NRPS families** |  |  |  |  |  |
| **BC- NRPS 1** | *Bacillus* . sp. BH32 | SJAS00000000.2.2.region002 | 286,950–346,871 | polyoxypeptin | **5%** |
| **BC- NRPS 2** | *Bacillus* sp. BH32 | SJAS00000000.2.15.region001 | 61,643–91,585 | - | - |
| **BC- Bacillibactin-like** | *B. cereus* A1 | NZ_CP015727.1.region003 | 1,501,835–1,550,463 | bacillibactin | **46%** |
| **BC- NRPS-like 1** | *B. cereus* G9842 | NC_011772.1.region001 | 386,831–430,076 | - | - |
| **BC-NRPS 3** | *B. cereus* G9842 | NC_011772.1.region006 | 2,287,034–2,352,013 | - | - |
| **BC-NRPS 4** | *B. cereus* ZB201708 | NZ_CP030982.1.region003 | 791,630–836,873 | chejuenolide A / chejuenolide B | **7%** |
| **Terpene family** |  |  |  |  |  |
| **BC- terpene 1** | *B. thuringiensis* L-7601 | NZ_CP020002.1.region012 | 3,779,510–3,801,363 | molybdenum cofactor | **17%** |
| **Siderophore families** |  |  |  |  |  |
| **BC- Petrobactin 1** | *B. cereus* G9842 | NC_011772.1.region004 | 1,852,404–1,866,111 | petrobactin | **100%** |
| **BC- Petrobactin 2** | *B. cereus* A1 | NZ_CP015727.1.region002 | 1,182,329–1,196,036 | petrobactin | **100%** |
| **Betalactone family** |  |  |  |  |  |
| **BC- Fengycin-like 1** | *B. cereus* G9842 | NC_011772.1.region007 | 2,387,411–2,412,649 | fengycin | **40%** |

**Fig. 4.**
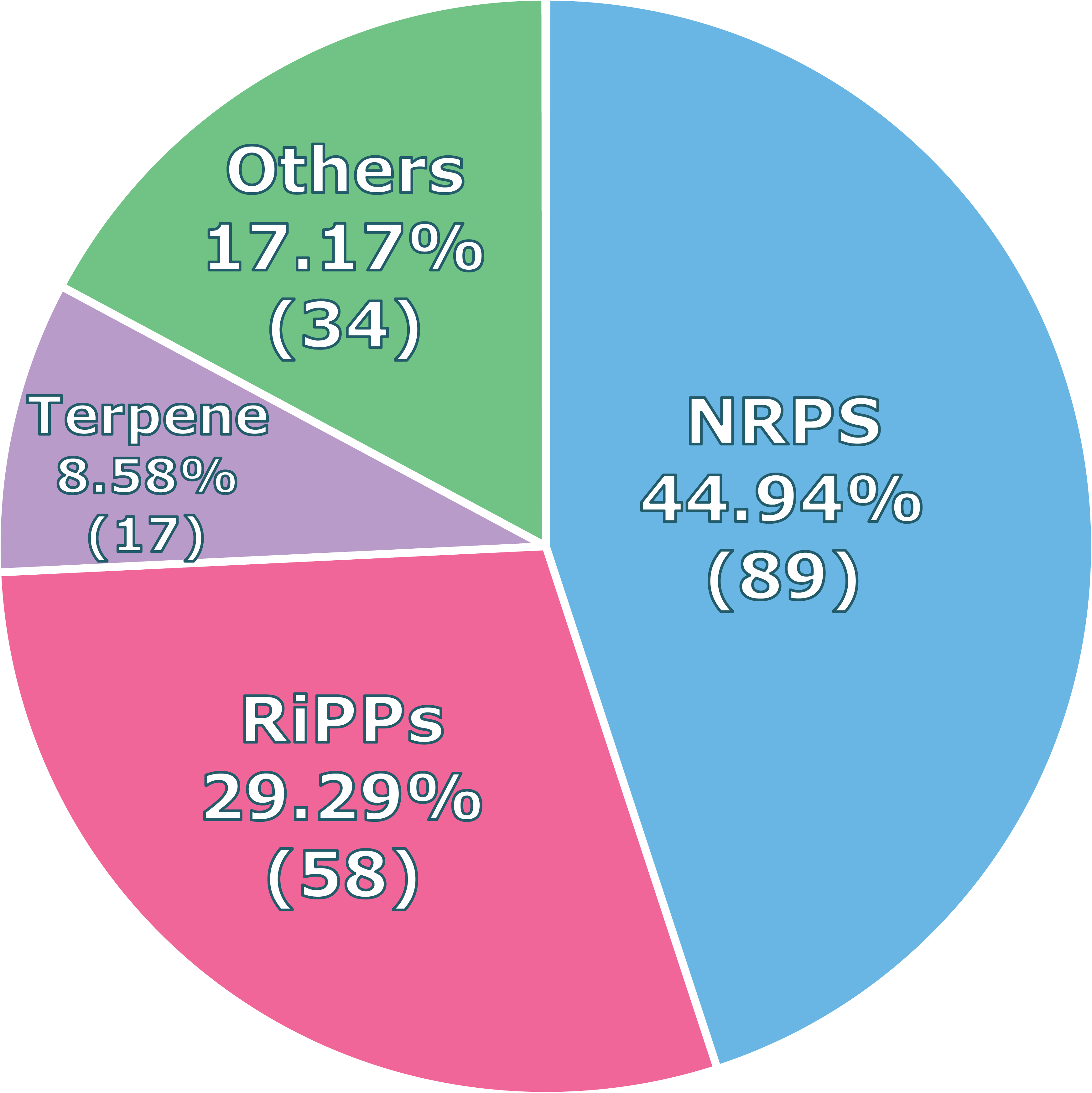
BGCs fractions (counts and percentages by class, according to BIG-SCAPE output).

Hence, we propose the following families:

#### The RiPPs gene cluster families of the *Bacillus cereus* group

The BIG-SCAPE output yielded 13 RiPPs GCFs (Fig. 5), with an average of 4 BGCs by family. It is the 2^nd^ largest group of BGCs with 58 clusters (including 7 singletons, table 4), without any known reference BGC in the MIBIG database (Fig. 5a). Following the proposed nomenclature, the identified families were named as follows (GCF name/ reference BGC strain and genomic region) (Fig. 5b; table 3; supp. Fig. S1):

• BC-Bacteriocin 1: *B. cereus* ZB201708, NZ_CP030982.1.region009
• BC-Bacteriocin 2: *B. thuringiensis* HD1002, NZ_CP009351.1.region005
• BC-Bacteriocin 3: *B. thuringiensis* serovar *galleriae* 4G5, NZ_CP010089.1.region007
• BC-Lasso peptide 1: *B. cereus* A1, NZ_CP015727.1.region001
• BC-Bacteriocin 4: *B. thuringiensis* serovar *morrisoni* BGSC 4AA1, NZ_CP010577.1.region009
• BC-Bacteriocin 5: *B. cereus* ZB201708, NZ_CP030982.1.region010
• BC-Cerecidin: *B. thuringiensis* YGd22-03, NZ_CP019230.1.region011

**Table 4.** Reference BGCs for RiPPs singletons

| RiPPs<br>singletons | strain | region | genomic<br>location<br>(nt) | most<br>similar<br>known<br>cluster | similarity |
| --- | --- | --- | --- | --- | --- |
| BCS-Bacteriocin 1 | *B. cereus* JHU | CP046511.1.region001 | 24,944 –<br>34,346 | - | - |
| BCS-Surfactin-like 1 | *B. cereus* ZB201708 | NZ_CP030982.1.region002 | 748,698 –<br>771,271 | - | - |
| BCS-Bacteriocin 2 | *B. cereus* JHU | CP046511.1.region002 | 128,885 –<br>139,342 | - | - |
| BCS-Bacteriocin 3 | *B. thuringiensis* c25 | NZ_CP022345.1.region010 | 4,376,042 –<br>4,388,249 | - | - |
| BCS-Bacteriocin 4 | *B. cereus* ZB201708 | NZ_CP030982.1.region014 | 5,325,429 –<br>5,338,733 | - | - |
| BCS-Bacteriocin 5 | *B. thuringiensis* L-7601 | NZ_CP020002.1.region013 | 5,353,484 –<br>5,365,667 | - | - |

Families BC-Bacteriocin 1, BC-Bacteriocin 2 and the singleton BC-Bacteriocin 3 are part of the RiPPs Clan I, while the families BC-Bacteriocin 4 and BC-Bacteriocin 5 are part of the RiPPs Clan II (Fig. 5c).

The BC-Cerecidin family was named based on the cerecidin compounds (lanthipeptides), with the closest known BGC (MIBIG database) being that of the cerecidin / cerecidin A1 / cerecidin A2 / cerecidin A3 / cerecidin A4 / cerecidin A5 / cerecidin A6 / cerecidin A7 (lanthipeptides putative class II), with a similarity of 94%.

The BC-Lasso peptide 1 was named based on its similarity with a LAP–bacteriocin BGC (sactipeptide/lassopeptide).

Meanwhile the singletons are (Fig. 5c, table 4; supp. Fig. S1):

• BCS-Bacteriocin 1: *B. cereus* JHU, CP046511.1.region001
• BCS-Surfactin-like 1: *B. cereus* ZB201708, NZ_CP030982.1.region002
• BCS-Bacteriocin 2: *B. cereus* JHU, CP046511.1.region002
• BCS-Bacteriocin 3: *B. thuringiensis* c25, NZ_CP022345.1.region010
• BCS-Bacteriocin 4: *B. cereus* ZB201708, NZ_CP030982.1.region014
• BCS-Bacteriocin 5: B. thuringiensis L-7601, NZ_CP020002.1.region013

The largest family was named BC-Lassopeptide 1 (with 17 analogous BGCs present in all of the studied genomes), followed by BC-Bacteriocin 4 (with 14 analogous BGCs). The remaining families have somewhere between 7 and 1 BGC; while the smallest family is composed of only 2 analogous BGCs (the BC-Cerecidin family) (Fig. 5b,c).

Furthermore, the RiPPs class is characterized by 7 singleton BGCs (Fig. 5c), displaying the highest number of singletons among the analyzed BGCs classes, which makes the RiPPs class the best niche for peculiar and likely to be unique secondary metabolites in the *B. cereus* group, in addition to pinpointing to possible horizontal gene transfer (HGT) at a BGC-level.

The singleton BCS-Surfactin-like 1 was named this way, due to its similarity (8%) with a known lanthipeptide BGC coding for the surfactin lipopeptide.

**Fig. 5.**
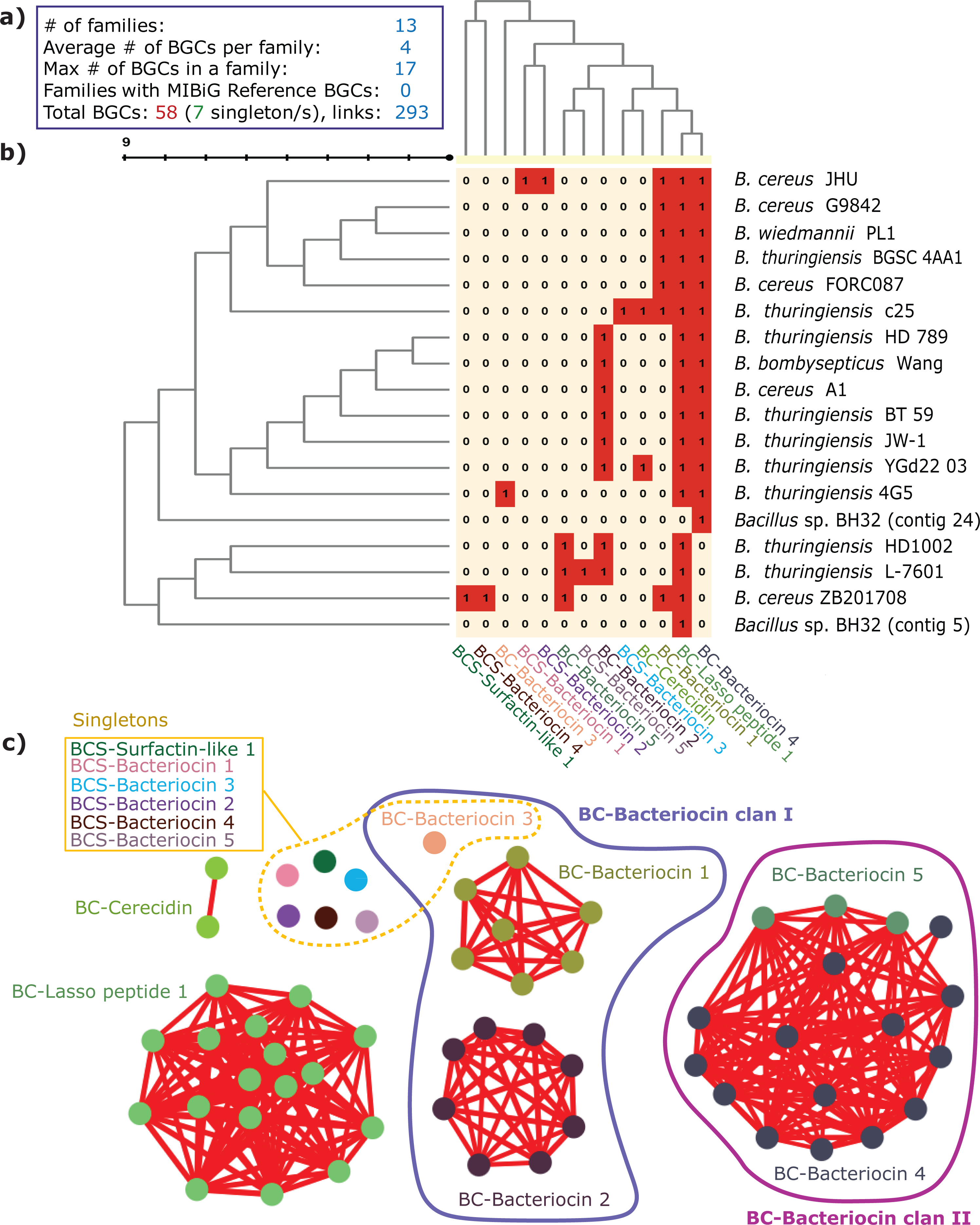
RiPPs BGC families/clans. **a)** RiPPs BGCs families statistics, **b)** proposed RiPPs families distribution matrix (presence: 1, absence: 0) among the *B. cereus* group genomes, **c)** RiPPs BGCs families/clans cluster networks (generated by the highest cutoff selected) and singletons (separate dots).

#### The NRPS gene cluster families of the *Bacillus cereus* group

For the NRPS class (Fig. 6), the BIG-SCAPE approach highlighted 6 families, without any reference BGC being known, which pinpoints further the future possible discovery of numerous novel metabolites in this intricate group. The NRPS class has the most represented families among the studied genomes, composed of 89 BGCs with an average number of 15 BGCs by family, and a staggering number of links (634 links) (Fig. 6a). No clans were proposed for this class.

We proposed the following names for the described NRPS families (Fig. 6b, table 3; supp. Fig. S2):

• BC-NRPS 1: *Bacillus.* sp. BH32, SJAS00000000.2.2.region002
• BC-NRPS 2: *Bacillus* sp. BH32, SJAS00000000.2.15.region001
• BC-Bacillibactin-like: *B. cereus* A1, NZ_CP015727.1.region003
• BC-NRPS-like 1: *B. cereus* G9842, NC_011772.1.region001
• BC-NRPS 3: *B. cereus* G9842, NC_011772.1.region006
• BC-NRPS 4: *B. cereus* ZB201708, NZ_CP030982.1.region003

The BC-NRPS 3 (showing low similarity with known BGCs coding for nostocyclopeptide A2: 28%, gramicidine: 16% from the MIBIG database; and the kurstakin C12 with a score of 0.639 according to the norine database) and the BC-Bacillibactin-like (showing a genes similarity of 46% with the known Bacillibactin BGC from the MIBIG database) families are represented in all of the studied genomes (17 analogous BGCs for both of them) (Fig. 6b,c).

It is noteworthy to mention that the BC-NRPS 4 family has a reference BGC showing little similarity with the known BGCs coding for polyketides chejuenolide A / chejuenolide B (7%) from the MIBIG database (table 3; supp. Fig. S2).

**Fig. 6.**
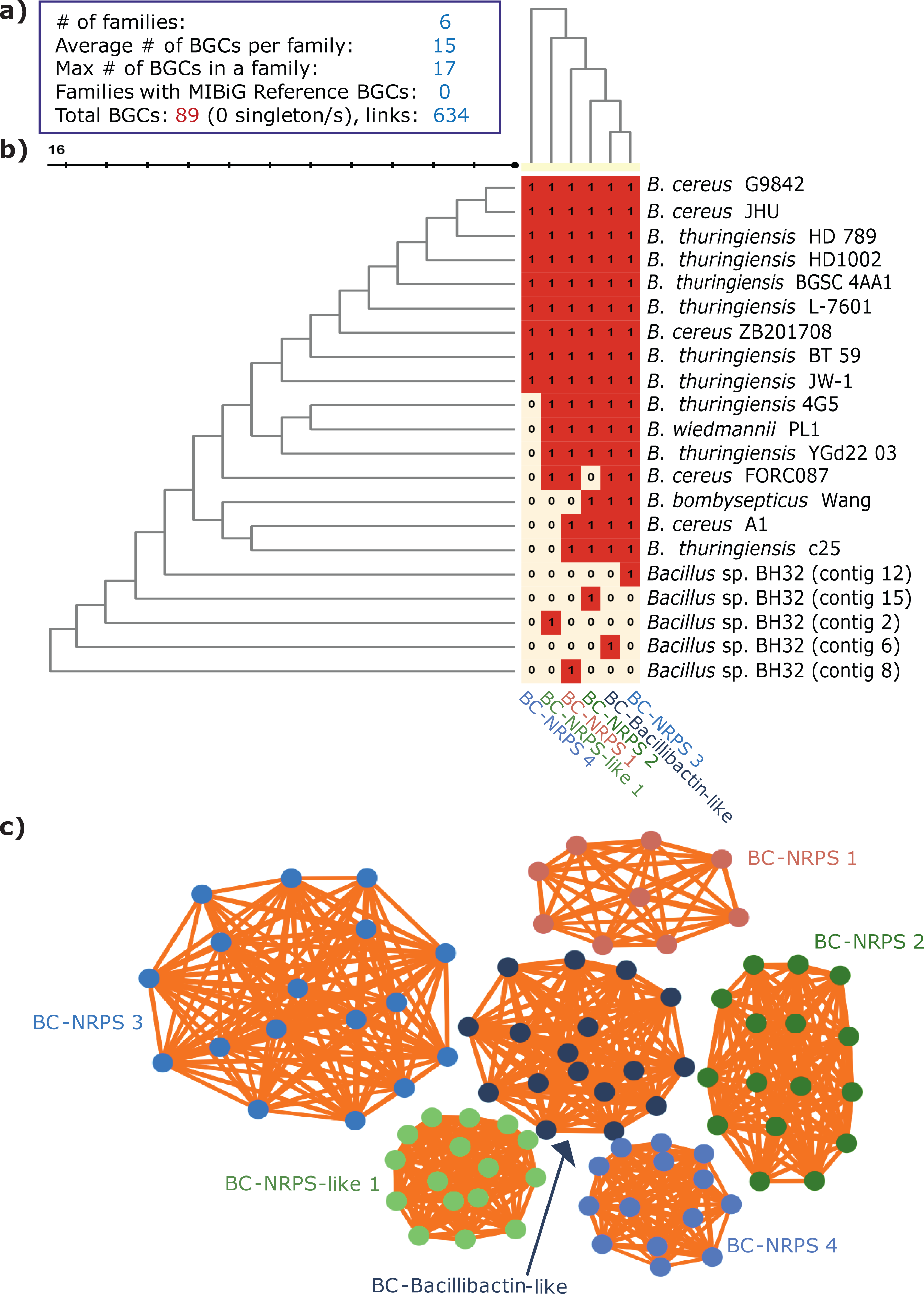
NRPS BGCs families. **a)** NRPS BGCs families statistics, **b)** proposed NRPS families distribution matrix (presence: 1, absence: 0) among the *B. cereus* group genomes, **c)** NRPS BGCs families cluster networks (generated by the highest cutoff selected).

#### The terpene gene cluster family of the *Bacillus cereus* group

For the terpene BGCs (Fig. 7), they all belong to the same family, named BC-Terpene 1, which bears similarity with the molybdenum cofactor (reference BGC: NZ_CP020002.1.region012 from *B. thuringiensis* L-7601, with a similarity of 17% with the molybdenum cofactor from the MIBIG database) (table 3; supp. Fig. S3).

It is the most conserved GCF (137 link in the corresponding similarity network) (Fig. 7 b) among the studied *B. cereus* genomes. The phylogenetic analysis (Fig. 7 c) confirms this conservation, showing two almost identical clades, with the exception of two different genes bearing 2 different pfam domains: the PF08445 FR47-like protein domain (Fig. 7 c, clade I) and the PF00583 Acetyl transferase GNAT family domain (Fig. 7 c, clade II).

**Fig. 7.**
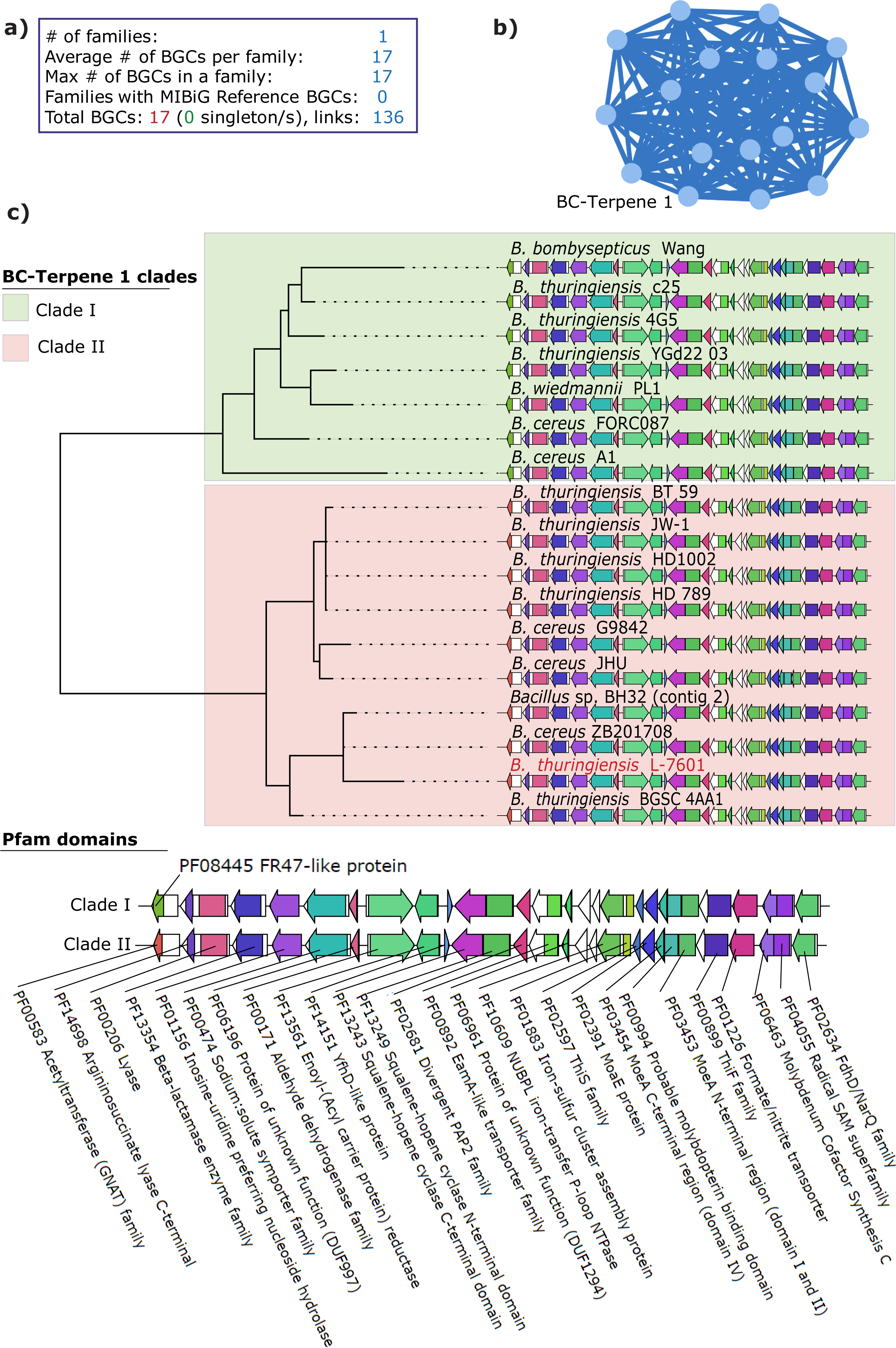
BC-Terpene 1 family features. **a)** BC-Terpene 1 BGCs family statistics, **b)** BC-Terpene 1 BGCs cluster network generated by the highest cutoff selected, **c)** CORASON-like tree generated for the BC-Terpene 1 GCF. This tree was created using the sequences of the Core Domains in the BC-Terpene 1 gene cluster family. These are defined as the domain type(s) that (1) appeared with the highest frequency in the BC-Terpene 1 gene cluster family and (2) were detected in the exemplar cluster (defined by the affinity propagation cluster), which is, in this case, that of *B. thuringiensis* L-7601 (in red). All copies of the Core Domains in the exemplar were automatically concatenated, as well as those from the best-matching domains of the rest of the BGCs in the BC-Terpene 1 gene cluster family (aligned domain sequences were used). The tree was inferred using FastTree43 with default parameters. Visual alignment was attempted based on the position of the ‘longest common information’ from the distance calculation step (between the exemplar BGCs vs. each of the remaining clusters). The Pfam domains of the 2 main clades are reported at the bottom part of this figure. The Pfam domains’ color significance can be retraced from a list, available here: https://git.wageningenur.nl/medema-group/BiG-SCAPE/blob/master/domains_color_file.tsv

#### The remaining siderophores and betalactone gene cluster families of the *Bacillus cereus* group

Regarding the remaining BGC classes (the 3 GCFs gathered under the section “others” in the BIG-SCAPE output)(Fig. 8), namely the BC-Petrobactin 1, BC-Petrobactin 2 families (both part of the BC-Petrobactin clan, coding for petrobactin siderophores, with a genes similarity of 100% with the known petrobactin BGC) and the BC-Fengycin-like 1 family (coding for betalactone products, with a genes similarity of 40% with known fengycin BGC from the MIBIG database), are another example of well-conserved families across the *B. cereus* group genomes, with an average number of BGCs by family of 11 (Fig. 8a).

The proposed families and their respective reference BGCs are (Fig. 8, table 3; supp. Fig. S4):

• BC-Petrobactin 2: *B. cereus* A1, NZ_CP015727.1.region002
• BC-Petrobactin 1: *Bacillus* sp. BH32, SJAS00000000.2.3.region001
• BC-Fengycin-like 1: *B. cereus* G9842, NC_011772.1.region007

Indeed, for the BC-Petrobactin clan (Fig. 8c), it has analogous BGCs across all of the studied genomes, being part of either the BC-Petrobactin 1 or BC-Petrobactin 2 families; and the BC-Fengycin-like 1 is represented in all of the 17 genomes (Fig. 8b), which is in line with the broad prevalence of Fengycin-like compounds reported in numerous strains belonging to the *B. cereus* group and their closely related taxa [38].

**Fig. 8.**
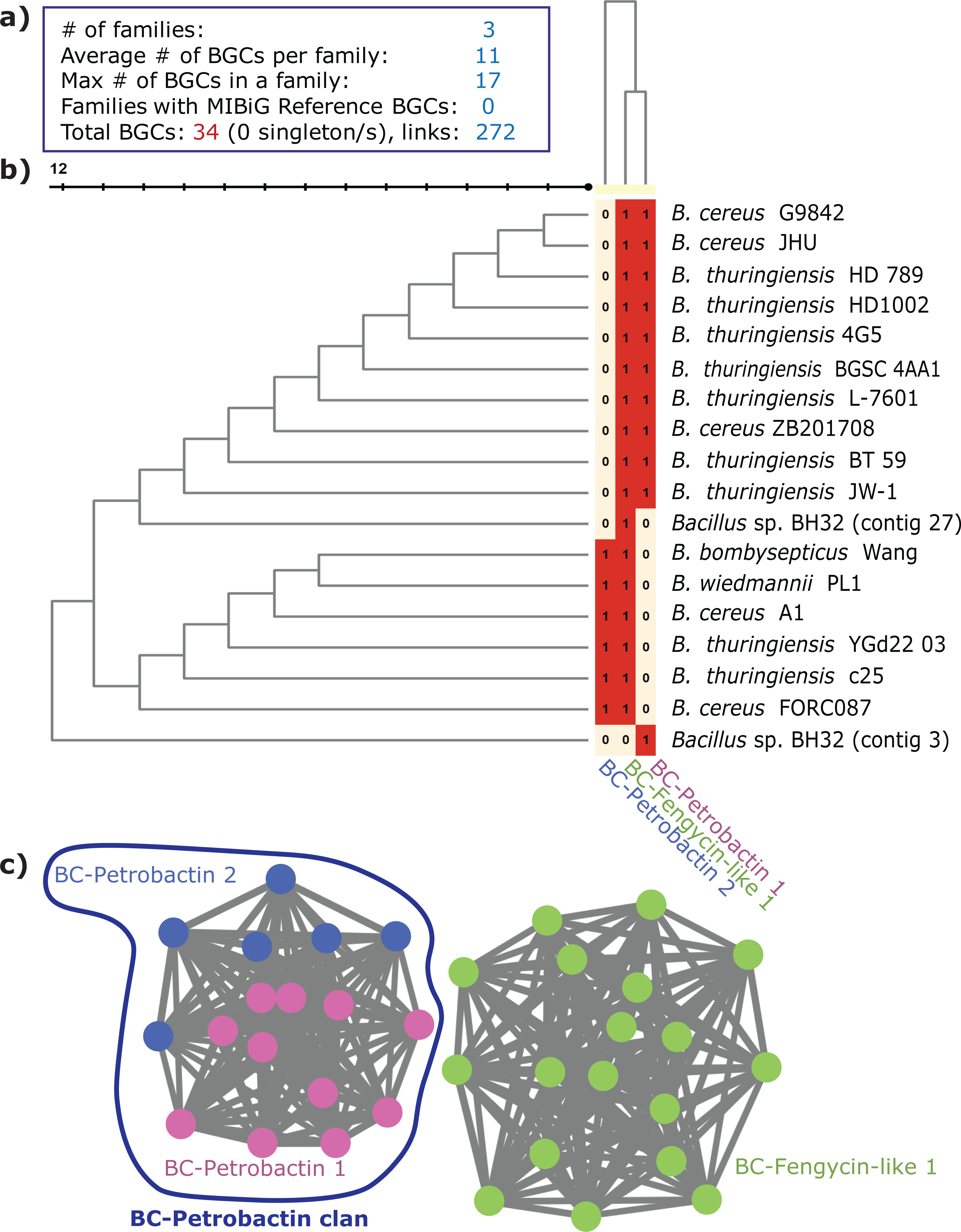
BC-Petrobactin 1, BC-Petrobactin 2, and BC-Fengycin BGCs families/clan. **a)** the remaining BGCs families statistics, **b)** proposed BC-Petrobactin 1, BC-Petrobactin 2, and BC-Fengycin families distribution matrix (presence: 1, absence: 0) among the *B. cereus* group genomes, **c)** BC-Petrobactin 1, BC-Petrobactin 2 and BC-Fengycin BGCs families/clan cluster networks (generated by the highest cutoff selected).

### 5. Harmony between the manual and the automatic approach

Based on the manual approach **(Fig. 3)**, we highlighted 2 BGCs synteny blocks “*synteny block A”* and “*synteny block B*”.

After performing a BIG-SCAPE analysis of the antiSMASH profiles of the studied genomes (Fig. 5, 6, 7, and 8), we could link the BGCs of the highlighted synteny blocks with their corresponding proposed families as follow:

For the **synteny block A:**

• BGC3 (bacteriocin: unknown) belongs to the **BC-Bacteriocin 4** family;
• BGC4 (bacteriocin: unknown) belongs to the **BC-Bacteriocin 1-2-3** families (BC-Bacteriocin clan I);
• BGC5 (betalacton, fengycin) belongs to the **BC-Fengycin-like 1** family;
• BGC6 (NRPS: gramicidin in *B. cereus* JHU; nostopeptolide A2 in *B. thuringiensis* serovar galleriae 4G5 ; unknown in the remaining strains) belongs to the **BC-NRPS 3** family;
• BGC7 (NRPS: bacillibactin) belongs to the **BC-Bacillibactin-like** family;
• BGC8 (siderophore: petrobactin) belongs to the **BC-Petrobactin** clan; and
• BGC9 (linear azol(in)e-containing peptides “LAP” bacteriocin) belongs to the **BC-Lasso peptide 1** family.

For the second conserved group named **“*synteny block B”***:

• BGC1 (terpene: molybdenum co-factor) belongs to the **BC-Terpene 1** family; and
• BGC2 (NRPS: polyoxypeptin, except for *B. thringiensis* c25, unknown) belongs to the **BC-NRPS 1** family.

That is to say, the synteny block A is a series of BGCs belonging (in order) to the families /clans (with the corresponding number of analogous BGCs found in each family):

BC-Bacteriocin 4 (14 BGCs), BC-Bacteriocin clan I (7+8+1 BGCs of the BC-Bacteriocin 1-2-3 families, respectively), BC-Fengycin-like 1 (17 BGCs), BC-NRPS 3 (17 BGCs), BC-Bacillibactin-like (17 BGCs), BC-Petrobactin clan (families 1 and 2, 17 BGCs), BC-Lasso peptide 1 (17 BGCs);

And the synteny block B is composed of BGCs belonging (in order) to the families: BC-Terpene 1 (17 BGCs), and BC-NRPS 1 (16 BGCs).

The remaining families are either under-represented across the *B. cereus* group or considered as singletons, which confirms the harmony and complementarities between both manual and automated approaches with BIG-SCAPE.

## Discussion

### 1. The importance of BGCs investigations

Deciphering beneficial features in plant growth-promoting bacteria requires research into the encoded parvome (the secondary metabolome inferred from the genome) [39]. Genes responsible for the production of secondary metabolites (SMs) are typically grouped into often quite large and complex BGCs [40]. BGCs are self-contained sets of co-located genes that accomplish the coordinated and regulated biosynthesis of a single set of SM congeners, with some exceptions, including BGCs that lack genes for required modifying enzymes that are located in different parts of the genome; distributed BGCs where two or more sub- clusters located in different parts of the genome collaborate during convergent biosynthesis of a single set of SM congeners; and superclusters that contain intertwined genes for the biosynthesis of more than one set of SM scaffolds [39, 41, 42].

BGCs are organized around genes that encode biosynthetic enzymes that yield the SM carbon skeleton (“backbone” enzymes), such as NRPSs, PKSs, PKS–NRPS hybrids, etc. The BGCs also feature genes encoding various enzymes that further modify the SM carbon skeleton (such as cytochrome P450 monooxygenases, various other oxidoreductases, etc) together with genes for transporters, regulators, and self-resistance determinants. In addition, many BGCs also harbor genes for enzymes that synthesize specialized monomers for the corresponding pathway [43].

### 2. BGCs screening and mining approaches

A biological approach in BGCs screening is considered the best prospect for resolving the potential of microbial parvomes. However, **Baltz** [44] estimated that 10^7^ strains would need to be examined to discover a novel class of antibiotics [44]. Thus platforms combining different approaches are the appropriate strategy [45]. Moreover, the apparent failure to uncover the full potential of natural product-producing microorganisms is likely due to the lack of understanding that is required to activate the expression of their BGCs in the laboratory [46]. Hence, new strategies for microbial SMs discovery are being employed, comprising genomics, metabolomics, and analytical tools that permit the study of complex systems [47].

Advances in genomics have unveiled a vast reservoir of BGCs in microbial genomes [46]. There are two main approaches to predicting BGCs. The first is based on pre-computed pHMMs derived from a set of genes known to participate in SM metabolism to identify sequences of interest [48, 49]. The second uses some function-agnostic criteria, such as synteny conservation or shared evolutionary history, to implicate genes as part of a gene cluster [50] which highlights the importance of synteny analysis among BGCs. Moreover, due to common metabolic functions across distant taxa, approaches employed by SMURF and antiSMASH are enormously successful [51]. Given the size of genomic data, studying BGCs on a case-by-case basis is no longer interesting. Hence, sequence similarity networking approaches can automatically relate predicted BGCs to gene clusters of known function and group them into gene cluster families (GCFs) [12, 46, 52, 53]. “Old school” methods gave way to new workflows that entail: (1) sequencing of the whole genome of strains that produce interesting SMs; (2) bioinformatic prediction of all BGCs; (3) comparison of the predicted BGCs with the retro-biosynthetic assessment of the structure of the target SM; (4) comparative genome analysis with organisms that produce similar SMs and with taxa that are phylogenetically close but not known to yield the target SM; and (5) comparative analysis of transcriptomes under SM producing vs. nonproducing conditions [39]. This led to the differentiation between BGCs that are likely to produce known compounds and BGCs that may encode novel chemistry. However, the number of GCFs to which no known functions can be linked is so great that it is difficult to know which of the BGCs encode the most interesting molecules [46].

### 3. Original BGC exploration methods

Out-of-the-box approaches are encouraged in targeting underexplored environmental niches and bacterial phyla [54]. Rare environments have rendered an interesting source of SMs. The chemical diversity of natural products correlates with the diversity of source microorganisms. This is probably due to the evolution of organism-specific biosynthetic machinery selected based on the adaptation of the microbe to the habitat, where beneficial SMs play an important part [47]. Applying an isolate-based genome-mining approach on bacteria obtained from unusual environments -such as deserts and arid regions like the sampling sites of the present study-can be used for screening unique BGCs that may govern the biosynthesis of novel natural products [54].

### 4. Automated bioinformatics pipelines vs. manual bioinformatics

Automated bioinformatics pipelines and manual bioinformatics are both methods used to analyze biological data, but they have some key differences. Automated bioinformatics pipelines are computer programs that are designed to perform specific bioinformatics tasks, such as sequence alignment, gene annotation, and variant calling, in a pre-defined, step-by-step manner. They are typically used to analyze large amounts of data and can be run on high-performance computing clusters, allowing for efficient and rapid data processing [55].

Automated bioinformatics pipelines are also designed to be easy to use and can be run by researchers with minimal bioinformatics experience [56]. Manual bioinformatics, on the other hand, is the process of analyzing biological data using manual methods, such as web-based tools, command-line programs, or custom scripts [57]. This method is typically used for smaller data sets, or when an automated pipeline is not available or not suitable for the task at hand. Manual bioinformatics requires a higher level of bioinformatics expertise and can be more time-consuming, but it can also be more flexible and customizable [58]. Both automated bioinformatics pipelines and manual bioinformatics have their advantages and disadvantages. Automated pipelines can be faster and more efficient at processing large amounts of data, but they may not be as flexible or customizable as manual methods.

Manual bioinformatics requires more expertise and can be more time-consuming, but it can also be more tailored to specific research questions.

During our study, Big-scape has proven to be useful in the exploration of BGCs in the *B. cereus* group thanks to its ability to identify and analyze large numbers of BGCs comprehensively and efficiently.

As Big-scape allowed the comparison of BGCs across different strains of *Bacillus cereus*, it helped us identify conserved and divergent BGCs among the *B. cereus* group, which will ultimately provide insights into the evolution and adaptation of *Bacillus cereus* to different environments.

### 5. The lack of BGCs classification

Despite the importance of BGCs, they are often difficult to classify and identify due to their complex genetic organization and a large number of different types of BGCs that can be found in different bacterial species [9]. One of the main challenges in BGCs classification is the lack of consensus on the criteria and methods used to classify BGCs [59]. Different researchers may use different criteria, such as gene content, gene organization, or evolutionary relationships, to classify BGCs, which can lead to inconsistent and conflicting results. Another challenge is that many BGCs are not well-annotated, making it difficult to identify the genes and functions associated with each BGC [60]. This can be particularly difficult in the case of novel BGCs, which may not have been previously described in the literature. Additionally, the lack of a standardized nomenclature for BGCs can further complicate the classification process [52].

### 6. Highlighting the hidden BGC potential and synteny in the *B. cereus* group

Many BGCs are not accounted for in the corresponding parvomes, referred to as “orphan” BGCs. These are either ‘cryptic’ (cannot yet be linked to a product, activity, or phenotype) or ‘silent’ (the compound is known in another organism, but experimental validation is required) [46]. The development of facile strategies to induce the expression of these silent BGCs, and to assign SM structures to orphan clusters will allow us to elucidate the microbial parvomes [39], including those of the *Bacillus cereus* group.

### 7. Lack of data about BGCs conservation in *Bacillus cereus* sensu lato

There is currently limited data available about the conservation of biosynthetic gene clusters (BGCs) in the *Bacillus cereus* group [61]. One reason for this is that the *Bacillus cereus* group is a large and diverse group of bacteria, comprising several closely related species and subspecies. This diversity can make it challenging to identify and study BGCs across the different strains of *Bacillus cereus*. Additionally, the identification and characterization of BGCs often require sophisticated techniques such as genome sequencing, transcriptomics, and metabolomics, which can be time-consuming and expensive [62].

Therefore, not all strains of *Bacillus cereus* have been fully characterized in terms of their BGCs. Furthermore, the majority of studies focus on certain strains of *Bacillus cereus*, such as *Bacillus anthracis*, *Bacillus thuringiensis*, and *Bacillus cereus* sensu stricto, that are known to produce important natural products [63–69], leaving out other strains.

The availability of genomes allowed the functional validation of identified BGCs based on the structures of known SMs. However, comparative genomics can also be used to derive hypotheses for the structures of the products of orphan BGCs. Thus, *de novo* sequenced BGCs may be assigned to known SM structural families if the core genes and the constituent tailoring genes are orthologous (or even syntenic) to functionally characterized BGCs, as those revealed in the present work. The synteny and the tight phylogenetic distances observed in our study support the conclusion that the BGCs in the *B. cereus* group arose dependently with the acquisition of conserved core component genes.

In the case of the *B. cereus* group, we observed that many of the biosynthetic gene clusters responsible for the production of natural products are syntenic, meaning that they are located in the same chromosomal position and have a similar gene organization among different strains of *B. cereus*. This suggests that these biosynthetic gene clusters have been inherited through common ancestry, and have been conserved over time through selective pressures.

The synteny of these gene clusters can be used to help understand the evolutionary relationships among different strains of *B. cereus*, and can also aid in the identification of new natural products. The growing number of genomes in databases revealed lineage-specific conservation of certain orthologous BGCs [39], as for the synteny bock A and B of the *B. cereus* group highlighted by the present study, or by pointing out the absence of certain widely present BGCs in some species, thereby allowing the generation of evolutionary hypotheses correlating the production of a given SM with the lifestyle and the evolutionary history of the producer [39].

Additionally, the synteny of BGCs can also aid in the development of new antibiotics and other bioactive compounds [10]. By identifying conserved regions of BGCs among different strains of *B. cereus*, researchers can infer the presence of similar biosynthetic pathways and enzymes among these strains, and can then use this information to guide the development of new antibiotics and other bioactive compounds. Overall, highlighting the synteny of BGCs can provide a valuable tool for understanding the biology, evolution, and biotechnology of the *B. cereus* group [70].

For instance, a syntenic BGC with orthologous genes to those of the *B. bassiana* oosporein BGC has recently been identified in the genome of *Cordyceps cycadae*, and the production of oosporein was confirmed by HPLC [71], and comparative genomics of *C. militaris* and *A. nidulans* revealed a syntenic BGC with four orthologous genes each in these fungi [72].

### 8. The RiPPs in the *B. cereus* group

RiPPs are an important class of natural products produced by the *Bacillus cereus* group. We highlighted a consequent diversity among this BGC class, as it appeared to be the most diversified compared to the remaining classes, with 13 RiPPs GCFs, and 7 singletons. RiPPs have a wide range of biological activities, and they have potential applications in medicine, agriculture, and industry [73]. RiPPs have a wide range of structural diversity and chemical complexity, making them an attractive target for natural product discovery. Additionally, the lack of resistance to RiPPs amongst pathogenic bacteria makes them an attractive target for the development of new antibiotics. The study of RiPPs biosynthesis and the enzymes responsible for the ribosomal synthesis and post-translational modification of these peptides is an active area of research, which has the potential to lead to the discovery of new RiPPs with unique properties and the development of new methods for RiPPs production [74].

We suggested that some BGCs could be part of mobile elements (BGC11, from the manual approach). One common method of BGCs acquisition is through integrative and conjugative elements (ICEs). The prevalence of these ICEs seems to be partially dictated by their ecological background: bacteria originating from soil, plants, or aquatic environments contain a greater number of ICEs than species from other environments [75].

### 9. BGCs in endophytes

Endophytes developed a variety of ways to successfully colonize plants; subdue their immunity and their physiology as a nutrient source, and defend the plant host from pathogens and opportunists. BGCs were also reported to mediate crucial functions in plant colonization by beneficial endophytes. These functions are often facilitated by the vast array of SMs produced by root-associated bacteria, which play a key role in inter- and intra-species interactions [76, 77]. To date, a handful of studies have explored the diversity and composition of bacterial SM-encoding BGC in soil [78].

Bioactive metabolites mediate important ecological functions, which are as diverse as their chemical structures. Siderophores enhance iron uptake in environments where the bioavailability of iron is limited [79], pigments protect against ultraviolet radiation and have antioxidant activity [80] and compatible solutes protect against osmotic stress [81]. Besides, it is possible to find in nature several examples of mutualistic relationships that have coevolved whereby the microorganisms are actively cultured in exchange for producing bioactive small molecules [47].

For instance, NRPS and PKS BGCs are responsible for the synthesis of a wide array of siderophores, toxins, pigments, and antimicrobial compounds [82] that are believed to play a pivotal role in bacterial adaptation to soil and rhizosphere ecosystems, and in plant health and development [83]. However, little is known regarding the distribution of these gene families in the root microbiome, and their functional role in the complex community interactions in root ecosystems remains an enigma [84].

Studying the encoded secondary metabolome of endophytes will amplify our understanding of the multiple roles that SMs play in the biotic and abiotic interactions in plants, leading to the unveiling of natural products that can be used for various applications.

#### Conclusions

The classification of Biosynthetic Gene Clusters (BGCs) is an evolving field that has gained significant attention in recent years. While some initial efforts have been made to classify BGCs, the field is still relatively new and the classification methodologies are constantly being refined and improved [22, 85]. Our work is an objective proposal for a consistent and standardized approach to BGCs classification among the *Bacillus cereus* group, based on a reproducible strategy that can be extended to other taxa, allowing comparison and integration of data from different studies to expand the initial classification scheme that we proposed. The current investigation is a substantive contribution to the discovery and characterization of new natural products and biosynthetic pathways, based on BGCs analogy and synteny.

## Supplementary Information

The online version contains available supplementary material.

**Additional file 1.** Supplementary figures.

**Additional file 2.** antiSMASH profiles.

**Additional file 3.** BIG-SCAPE output files.

## Funding

This work didn’t receive any funding.

### Authors’ contributions

HAB conceived, designed the study; and analyzed the data. HAB and AY drafted the manuscript; HAB, AY, AZ, and AM reviewed the manuscript. All authors read and approved the final manuscript.

## Supporting information

antiSMASH profiles + BIG-SCAPE output

## Acknowledgments

We gratefully acknowledge the inspirational impact of Dr. Livio Antonielli. Furthermore, our warmest appreciation for 2019’s EMBO genomics and bioinformatics course organizers, with special thanks to Dr. Fredj Tekaia.

## Availability of data and materials

The datasets supporting the conclusions of this article are included within the article (and its additional files). The genomic dataset can be accessed through the corresponding accession numbers. The antiSMASH profiles and the BIG-SCAPE output are available as supplementary material, from which the reference BGCs can be fetched (see table 3).

## Declarations

### Ethics approval and consent to participate

Not applicable.

### Consent for publication

Not applicable.

---

**Fig. S1** RiPPs reference BGCs for each proposed family. For each reference BGC, from top to bottom: reference BGC info (from left to right are mentioned: the name of the proposed RiPPs family; the strain bearing the reference BGC; the genomic region; and the most similar known cluster with the similarity %); a depiction of the BGCs regions/distribution among the genome; and the BGC organization (from antiSMASH output).

**Fig. S2** NRPS reference BGCs for each proposed family. For each reference BGC, from top to bottom: reference BGC info (from left to right are mentioned: the name of the proposed NRPS family; the strain bearing the reference BGC; the genomic region; and the most similar known cluster with the similarity %); a depiction of the BGCs regions/distribution among the genome; and the BGC organization (from antiSMASH output).

**Fig. S3** BC-Terpene 1 reference BGC. From top to bottom: reference BGC info (from left to right are mentioned: the name of the proposed family; the strain bearing the reference BGC; the genomic region; and the most similar known cluster with the similarity %); a depiction of the BGCs regions/distribution among the genome; and the BGC organization (from antiSMASH output).

**Fig. S4** Siderophore/Betalactone reference BGCs for each proposed family. For each reference BGC, from top to bottom: reference BGC info (from left to right are mentioned: the name of the proposed family; the strain bearing the reference BGC; the genomic region; and the most similar known cluster with the similarity %); a depiction of the BGCs regions/distribution among the genome; and the BGC organization (from antiSMASH output).

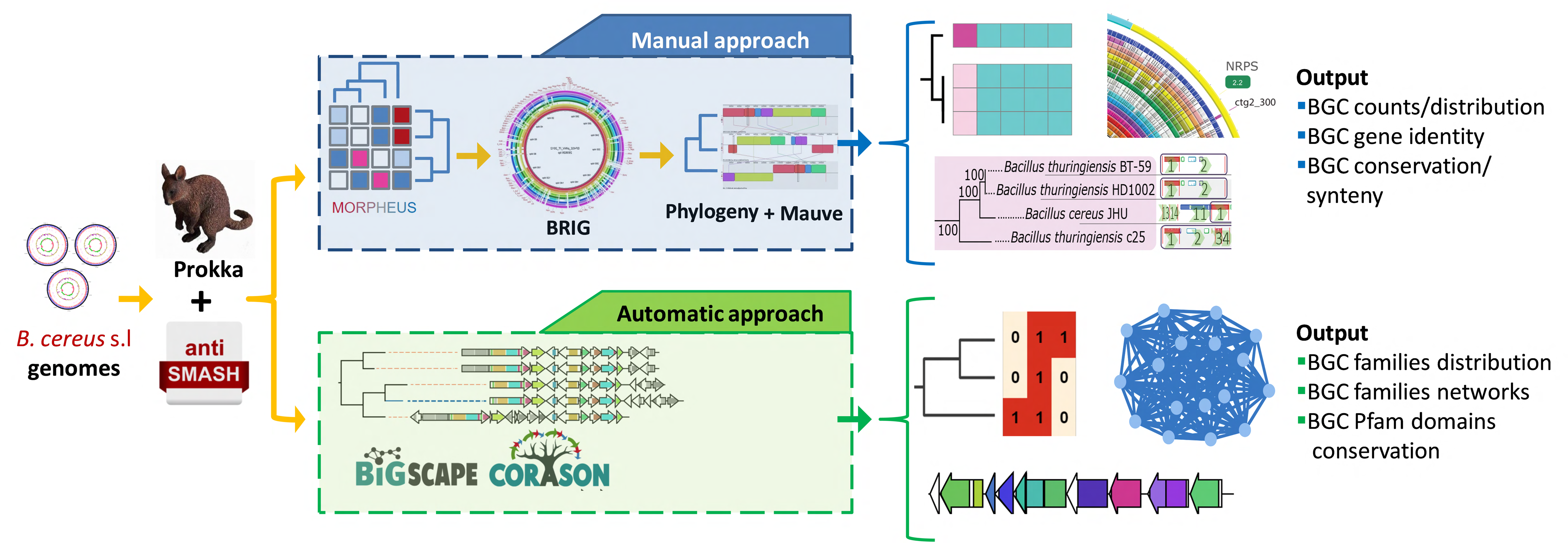

