## Supplementary figures and images for "Delving into the *Bacillus cereus* group biosynthetic gene clusters cosmos: a comparative-genomics-based classification framework"

### Supplemental Figure S1

#### RiPPs reference BGCs

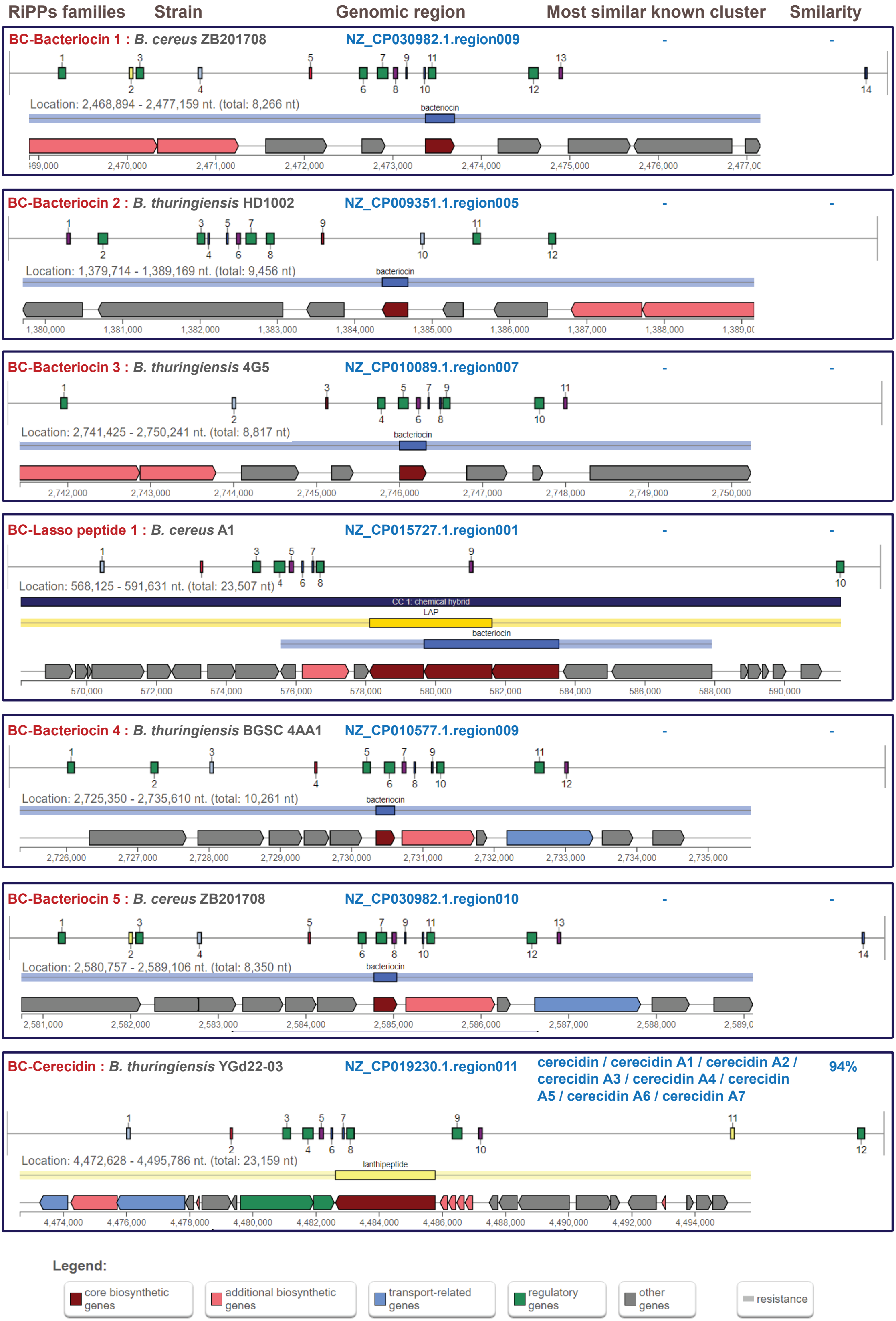

### Supplemental Figure S2

#### NRPS reference BGCs

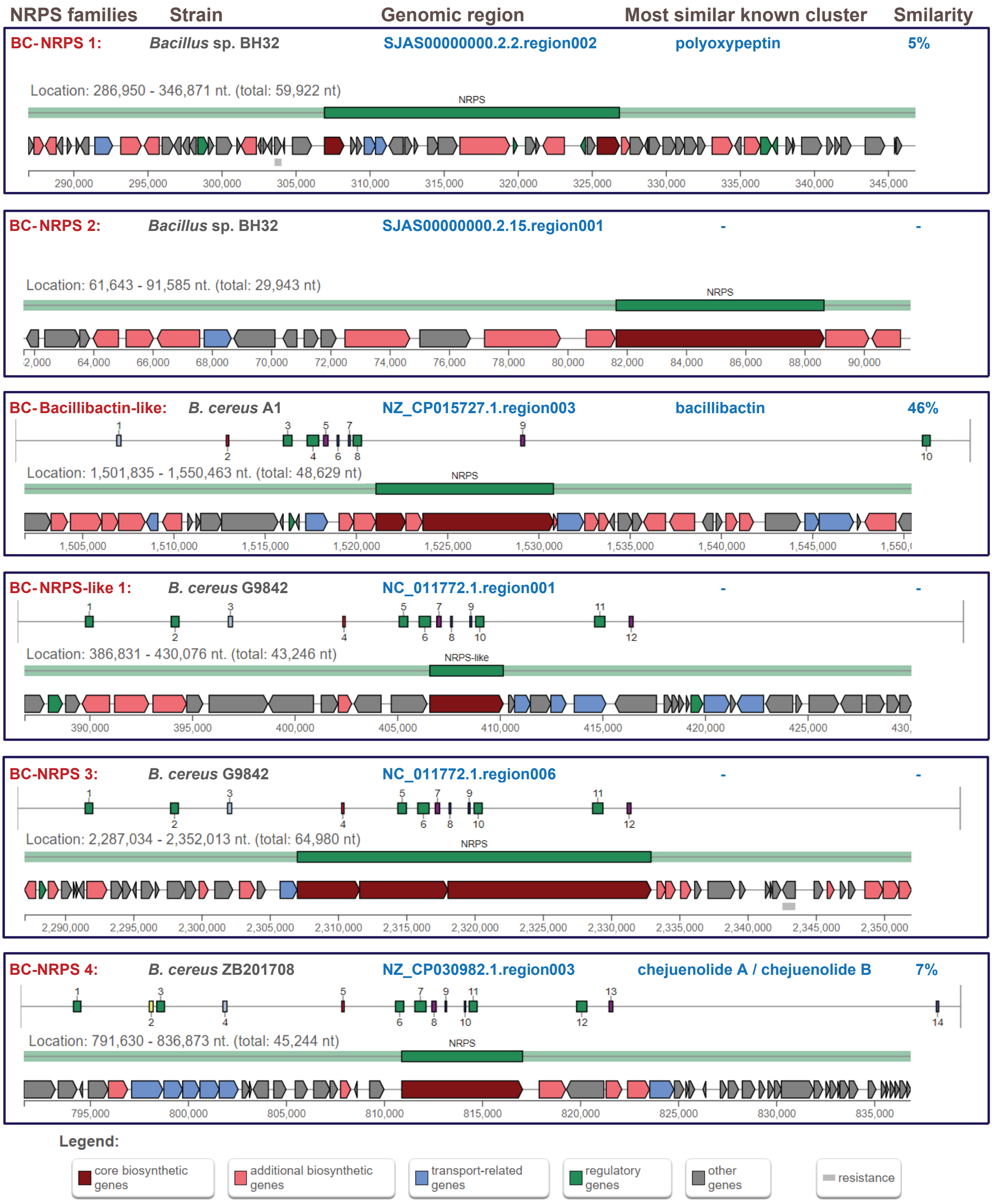

### Supplemental Figure S4

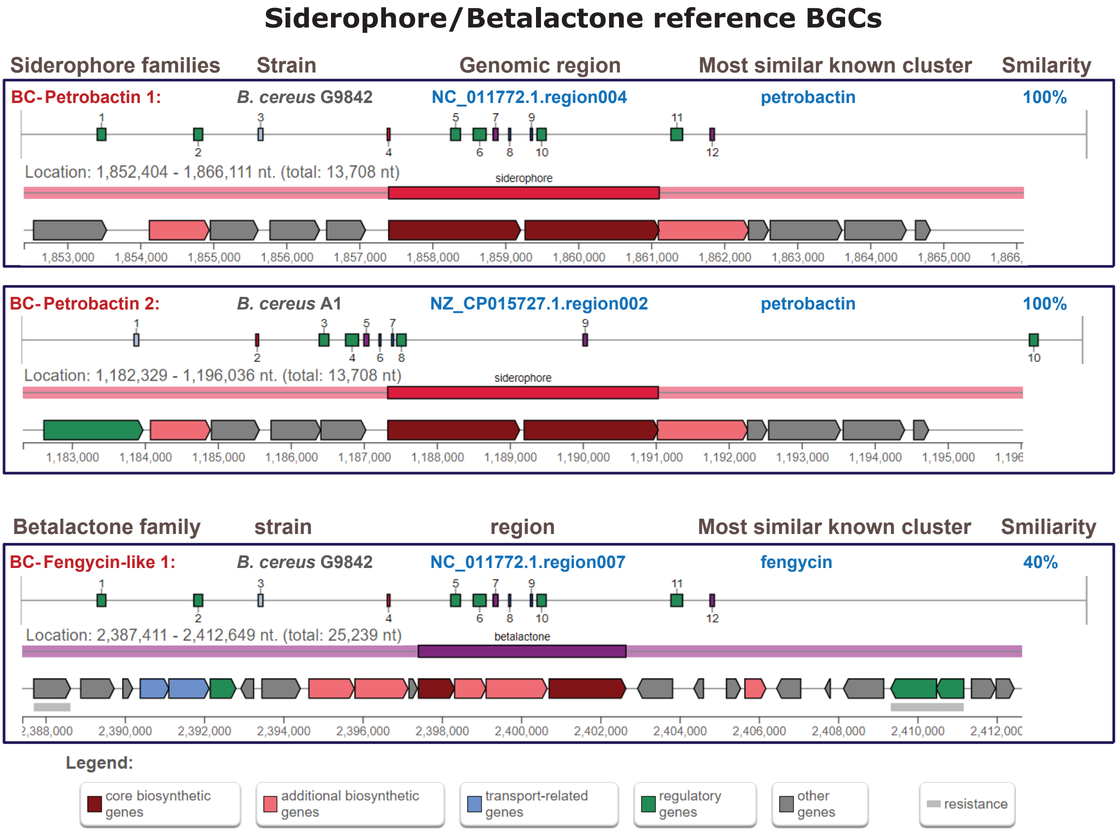
