## Supplemental Figure S3 for "Delving into the *Bacillus cereus* group biosynthetic gene clusters cosmos: a comparative-genomics-based classification framework"

### Terpene reference BGC

| Terpene family | Strain | Genomic region | Most similar known cluster | Smilarity |
|---|---|---|---|---|
| BC-terpene 1: | *B. thuringiensis* L-7601 | NZ_CP020002.1.region012 | molybdenum cofactor | 17% |

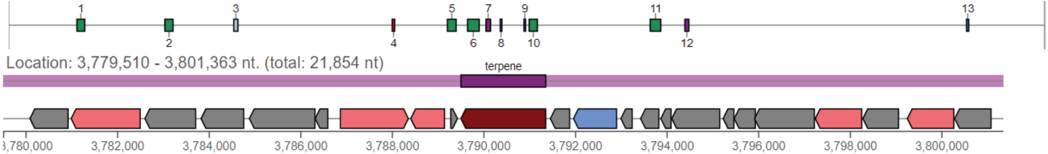


**Legend:**

- core biosynthetic genes
- additional biosynthetic genes
- transport-related genes
- regulatory genes
- other genes
- resistance
